# Rapid degradation and 3D CLEM of condensin reveal chromatin compaction uncoupled from chromosome architecture in mitosis

**DOI:** 10.1101/173633

**Authors:** Kumiko Samejima, Daniel G. Booth, Hiromi Ogawa, James R. Paulson, Linfeng Xie, Cara A. Watson, Melpomeni Platani, Masato T. Kanemaki, William C. Earnshaw

## Abstract

The requirement for condensin in chromosome formation in somatic cells remains unclear as imperfectly condensed chromosomes do form in conventional condensin-depleted cells. Here we have dissected the role of condensin at different stages of vertebrate mitosis by combining auxin-mediated rapid depletion of condensin subunit SMC2 with chemical genetics to obtain near-synchronous mitotic entry of chicken DT40 cells. We analysed the outcomes by live and fixed-cell microscopy methods, including 3D correlative light and serial block face scanning electron microscopy. Following rapid depletion of condensin, chromosomal defects were obvious. The chromatin was compacted normally, but formed a single mass of mitotic chromosomes clustered at one side of a bent mitotic spindle. Cultures arrest at prometaphase, eventually exiting mitosis without segregating chromosomes. Experiments titrating the auxin concentration suggest a previously unsuspected dual role of condensin, as different condensin levels are required for anaphase chromosome segregation and formation of a normal chromosome architecture.

## INTRODUCTION

As cells enter mitosis, the chromatin is compacted 2-3-fold in volume as it undergoes a dramatic re-organisation (Lleres et al., 2009; Martin and Cardoso, 2010; Vagnarelli and Earnshaw, 2012). Rod-shaped mitotic chromosomes form and chromosome scaffold proteins are localised along the sister chromatid axes (Paulson and Laemmli, 1977; Lewis and Laemmli, 1982; Earnshaw et al., 1985; Gasser et al., 1986; Saitoh et al., 1994; Samejima et al., 2012). These proteins include topoisomerase IIα, chromokinesin KIF4 and SMC2 (Structural Maintenance of Chromosomes 2), a core subunit of both condensin complexes (Earnshaw et al., 1985; Gasser and Laemmli, 1986; Saitoh et al., 1994; Hirano and Mitchison, 1994; Hirano et al., 1997; Ono et al., 2003; Samejima et al., 2012).

Loss of condensin leads to chromosome mis-segregation with masses of lagging chromosomes blocking cytokinesis in all the experimental systems tested including yeasts, *D. melanogaster*, nematodes, and vertebrate cells (Saka et al., 1994; Steffensen et al., 2001; Hagstrom et al., 2002; Hudson et al., 2003) (Houlard et al., 2015). Mitotic chromosome structural defects are also commonly observed. However, the extent of those defects varies between different experimental systems (Hirano and Mitchison, 1994; Hirano et al., 1997; Saka et al., 1994; Bhat et al., 1996; Steffensen et al., 2001; Hagstrom et al., 2002; Somma et al., 2003; Hudson et al., 2003; Coelho et al., 2003; Vagnarelli et al., 2006; Maddox et al., 2006; Samoshkin et al., 2009). At one extreme, immuno-depletion of condensins from *Xenopus* egg extracts results in the formation of a decondensed chromosome mass, indicating that condensins are essential for assembly and maintenance of mitotic chromosome structure (Hirano and Mitchison, 1994; Hirano et al., 1997). *S. pombe* temperature-sensitive mutants of condensin subunits also fail to condense chromosomes under restrictive conditions (Saka et al., 1994; Sutani et al., 1999). Meiotic I oocytes depleted of condensin subunits showed massively deformed chromosomes (Houlard 2015).

At the other extreme, vertebrate cells depleted of condensins using conditional knockouts or siRNA exhibit relatively moderate defects (Hudson et al., 2003; Ono et al., 2003; Ono et al., 2004; Vagnarelli et al., 2006; Samoshkin et al., 2009). Individual chromosomes are seen, but they are wider and appear to lack the structural rigidity seen in wild type chromosomes. These inconsistent phenotypes among different experimental systems pose a “condensin paradox” (Gassmann et al., 2004), suggesting that condensin might not be universally required for mitotic chromosome formation.

In the *Xenopus* egg extract experiments, condensin was immuno-depleted before the addition of sperm DNA. In the case of *S. pombe*, condensin was rapidly inactivated by a temperature shift. In contrast, vertebrate condensin was gradually lost by natural turnover over more than one cell cycle after synthesis of new protein was halted. Thus, the milder defects in chromosome assembly seen in vertebrates might be explained by cellular adaptation to the gradual loss and/or incomplete depletion of condensin [see e.g. (Wood et al., 2016)]. Furthermore, the various mitotic defects observed could result from a mixture of primary and secondary effects or even non-mitotic functions of condensin (Hirano, 2016). Together, these observations suggested that rapid depletion of condensin in vertebrate cells might more accurately reveal its true mitotic function(s).

This rapid depletion is now possible using an <u>A</u>uxin <u>I</u>nducible <u>D</u>egron (AID) system (Nishimura et al., 2009; Kanemaki, 2013). The plant hormone auxin enhances the affinity of the plant-specific F-box protein OsTIR1 for the AID tag AtIAA17. In the presence of auxin, tagged target proteins become poly-ubiquitylated and are degraded rapidly via the ubiquitin-proteasome pathway. It can take as little as 1 h to deplete a target protein in vertebrate cells. Furthermore, the AID system allowed us to study cells partially depleted of condensin by titrating the amount of auxin (Nishimura 2009). This is difficult or impossible using the TEV protease cleavage mediated condensin inactivation system.

A fundamental difficulty with studying mitotic chromosome formation is that chromosome morphology changes on a minute-by-minute basis as cells enter mitosis. However, prophase cells comprise less than 1 % of the population in asynchronous cultures. Thus, cell synchrony protocols are needed. Importantly, commonly used nocodazole or colcemid treatments are not suitable for studying chromosome morphology, as both treatments delay cells in prometaphase and induce hyper-condensation of the chromosomes.

An alternative strategy is to synchronise cultures before they enter mitosis. Regulated CDK1 activity is essential for mitotic entry and progression. Manipulation of this activity can be used to enrich for mitotic cells (Nurse and Bissett, 1981); (Vassilev et al., 2006; Smith et al., 2011). To this end, an analogue sensitive cyclin dependent kinase 1 (CDK1^as^) system was established to specifically inhibit CDK1 activity (Hochegger et al., 2007). CDK1^as^ has a mutation in the ATP binding pocket allowing the binding of 1NMPP1, an ATP analogue with a bulky side chain (Bishop et al., 2000). Addition of 1NMPP1 blocks CDK1^as^ cells in late G2, and upon washout of the drug, the cells swiftly enter mitosis.

Here we have used rapid degradation of SMC2 in highly synchronized cultures coupled with 3D-CLEM (correlative light and electron microscopy) to probe the function of condensin in mitotic chromosome structure and segregation. We find that mitotic chromatin is compacted to the normal extent though the intrinsic chromosome architecture is faulty in condensin-depleted cells. We also find that differing levels of condensin are required for normal chromosome architecture, sister chromatid axis formation and chromosome segregation at anaphase.

## RESULTS

### SMC2-AID-GFP cell line to study the mitotic roles of condensin

Defining the roles of condensin in vertebrate cells during mitosis has been difficult due to relatively mild defects observed following depletion of condensin subunits in mitotic cells of numerous species, compared to defects seen *in vitro*, in yeasts, in meiotic mouse oocytes and in fly embryos. This could be explained by variable levels of residual condensin in different experimental systems or by the cells adapting to the gradual loss of condensin. We hypothesized that rapid depletion of a key condensin subunit might resolve this issue. In order to test this hypothesis, we decided to modify SMC2 (condensin I and II common subunit) to be rapidly depleted in chicken DT40 cells. This is because both condensin I and II have important roles in mitotic chromosome assembly in chicken DT40 cells similar to human cultured cells (Ono 2003, Green 2012) and also we could directly compare the defects with those of our chicken DT40 conditional SMC2 knockouts (SMC2^ON/OFF^) (Hudson 2003).

Taking advantage of an inducible degron (AID) system, we established the SMC2-AID-GFP cell line expressing a SMC2 cDNA fused with a minimal AID degron and GFP tag as the sole source of SMC2 together with the auxin-responsive F-box protein TIR1 (Figure 1A and see Materials and Methods). SMC2-AID-GFP cells proliferated well only in the absence of auxin (Figure 1B). In contrast, addition of auxin did not affect the growth rate of wild type cells. Mitotic index (MI), mitotic profile and cell cycle profile of the SMC2-AID-GFP cells were similar to the ones of wild type cells without auxin (Figure 1-figure supplement 1A, Figure 2-figure supplement 1C). SMC2-mAID-GFP was expressed at a level similar to the endogenous SMC2 of wild type cells (Figure 1C) and localised correctly along the axis of sister chromatids in living cells (Figure 1D). Prometaphase chromosomes of SMC-AID-GFP cells appeared indistinguishable from those of wild type cells in the absence of auxin (Figure 1G, Figure 1-figure supplement 1B).

**Figure 1.**
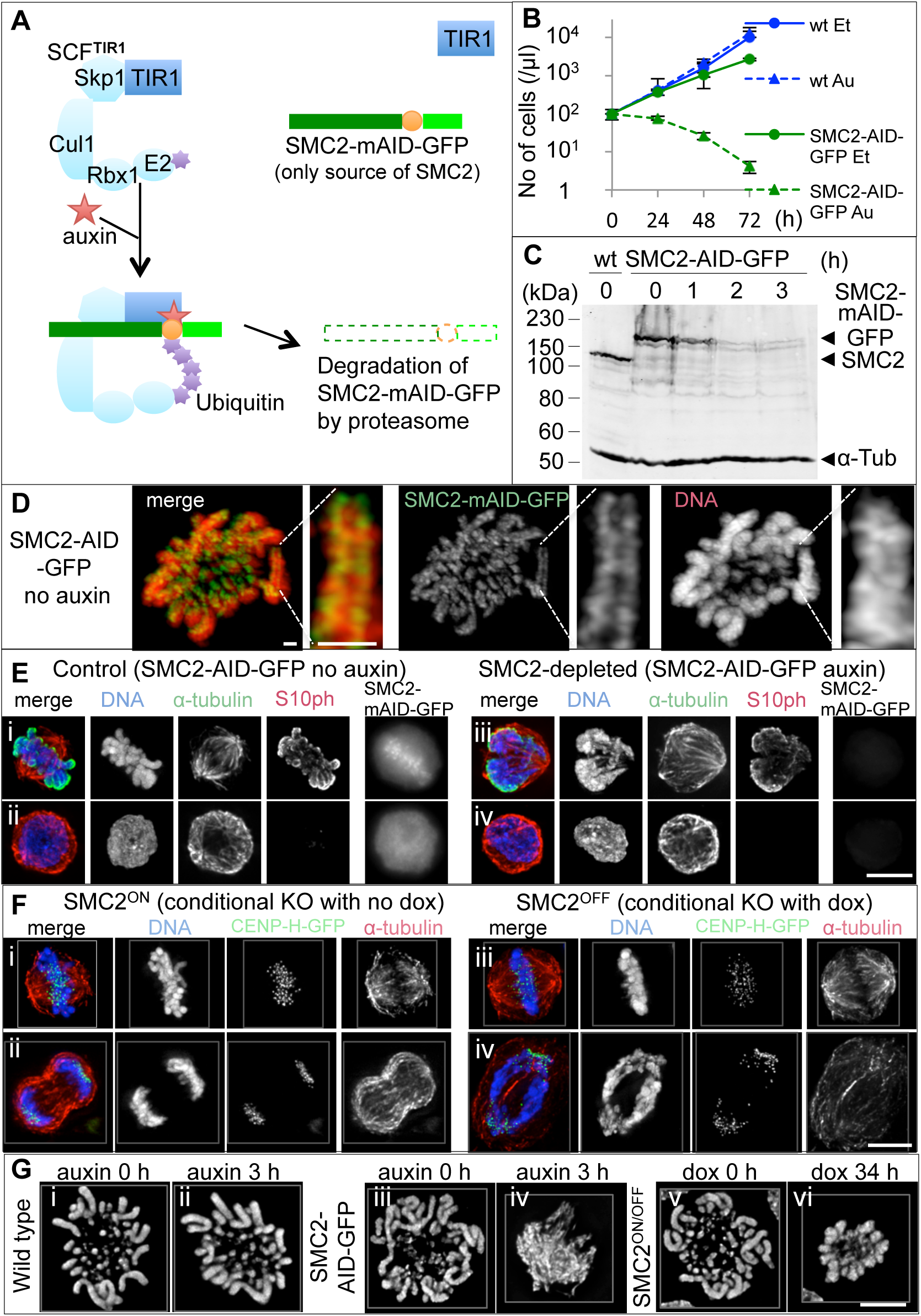
Rapid depletion of SMC2-mAID-GFP upon auxin treatment results in a severely defective mitotic chromosome formation. **(A)** Diagram introducing SMC2-AID-GFP cells. Auxin addition recruits the miniAID tag to the SCF^TIR1^ complex, resulting in rapid degradation of SMC2-mAID-GFP via the proteasome pathway. **(B)** Growth curve of wild type cells and SMC2-AID-GFP cells treated with either ethanol (Et: solvent) or 125 μM auxin (Au). Error bars show standard deviation. **(C)** Immunoblot analysis of asynchronous wild type and SMC2-AID-GFP cells. SMC2-AID-GFP cells were treated with 125 μM auxin for 0 – 3 h. SMC2 and SMC2-mAID-GFP were detected with anti-SMC2 antibody, α-tubulin was a loading control. **(D)** Live cell imaging of a SMC2-AID-GFP cell by Zeiss Airy microscope. DNA was stained with SiR-DNA. SMC2-mAID-GFP concentrated along the axis of sister chromatids. Scale bar 1 μm. **(E)** SMC2-AID-GFP cells treated with ethanol (solvent) (i, ii) or auxin for 3 h (iii, iv) were fixed with formaldehyde and stained for α-tubulin (green), and Histone 3 phospho-SerlO (red) and DNA (blue). The GFP signal was undetectable (iii, iv) and the shape of mitotic chromosomes (iii) was highly abnormal in SMC2-depleted cells. Scale bar 5 μm. **(F)** SMC2^ON/OFF^/CENP-H-GFP cells (Vagnarelli et al., 2006) converted to CDK1^as^ were treated with doxycycline for 0 or 30 h, stained in metaphase and anaphase for DNA (blue), CENP-H-GFP (green) and α-tubulin (red). **(G)** Wild type/ CDK1^as^ cells (0 or 3 h auxin treatment), SMC2-AID-GFP/CDK1^as^ cells (0 or 3 h auxin treatment) cells and SMC2^ON/OFF^/CDK1^as^ cells (0 or 34 h doxycycline treatment) were fixed with cold Methanol/Acetic acid and stained for DNA. More pictures in Figure1-figure supplement 1B. Scale bar 5 μm.

### Chromosomes condense into a single mass following rapid SMC2 depletion

Auxin addition induced the rapid depletion of SMC2-mAID-GFP in asynchronous cell cultures. After a 3 h auxin treatment, SMC2-mAID-GFP fell to approximately 5% of the wild-type level, the number of GFP-positive cells declined, and the GFP signal was lost (Figure 1C, E, Figure 1-figure supplement 1C). Chromatin of mitotic cells depleted of SMC2 clustered in a compact mass in which single chromosomes could not be resolved (Figure 1E-G, Figure 1-figure supplement 1B). The chromosome morphological defects seen in SMC2-AID-GFP cells treated with auxin were much more severe than those in SMC2^OFF^ cells treated with doxycycline for 30-34 h. SMC2^OFF^ (conventional conditional knockout) chromosomes were wider and fuzzier than controls, but individual chromosomes were still visible (Figure 1G, 2Ai, Figure 1-figure supplement 1B). Furthermore, SMC2^OFF^ cells underwent aberrant anaphase with chromosome bridges as previously reported (Figure 1Fiv). In previous reports, chromosome mis-segregation at anaphase was the most prominent phenotypic defect after condensin depletion or inactivation (Saka et al., 1994; Steffensen et al., 2001; Hagstrom et al., 2002; Hudson et al., 2003). In contrast, we were surprised to observe no anaphase or telophase cells following rapid SMC2 depletion (Figure 2C).

**Figure 2.**
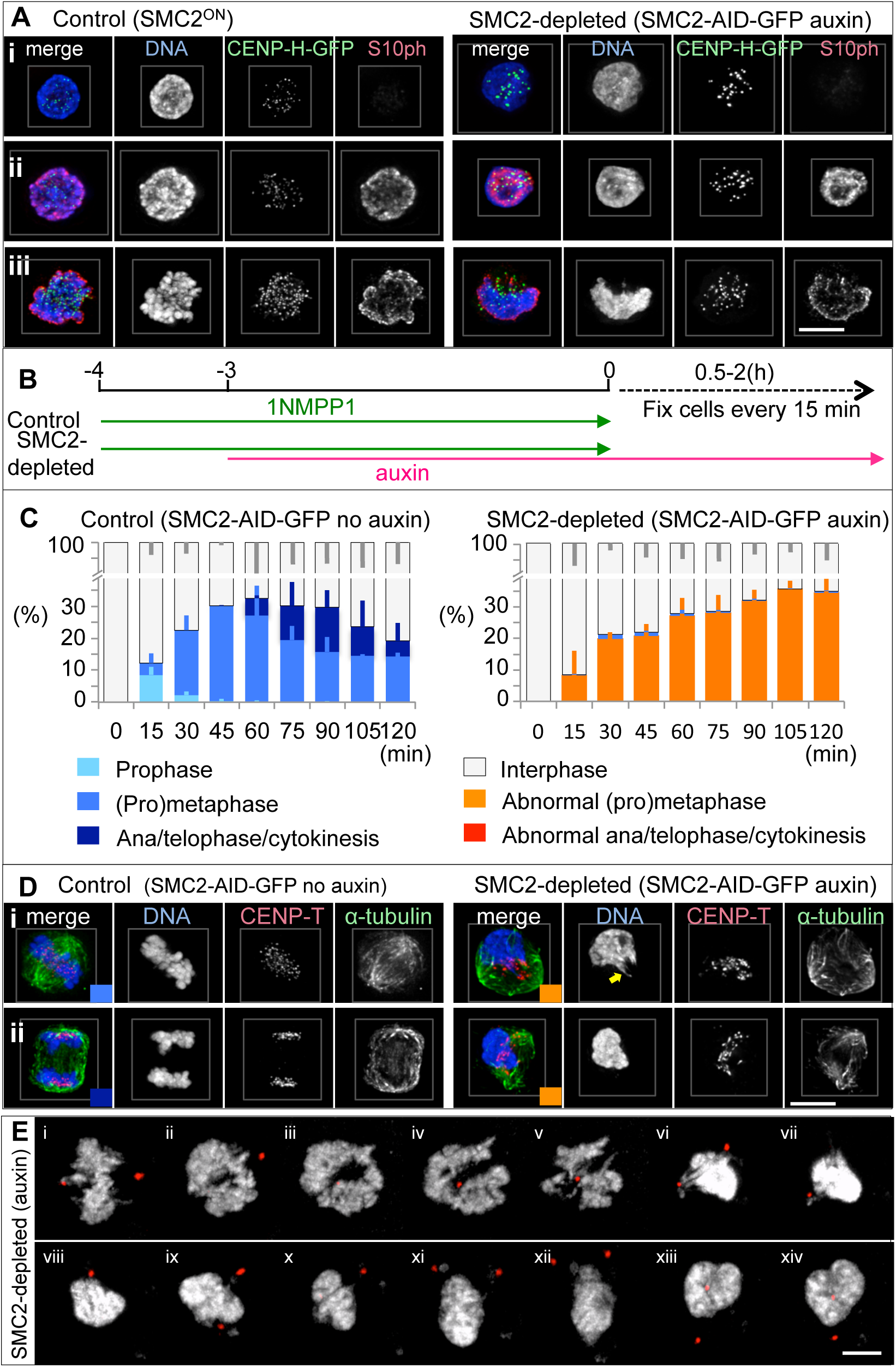
SMC2-depleted cells accumulate at prometaphase with a single chromosome mass. **(A)** Phosphorylation of Histone 3 SerinelO (red) was not affected by SMC2 depletion. DNA shown in blue and CENP-H-GFP in green. Interphase cells (i), prophase cells (ii) and prometaphase cells (iii). Control: SMC2^ON^/ CDK1^as^/CENP-H-GFP cells. SMC2-depleted: SMC2-AID-GFP/CDK1^as^/CENP-H-GFP cells. **(B)** Experimental timeline for auxin treatment. After 1NMPP1 washout, SMC2-AID/CDK1^as^ cells were fixed with 4 % formaldehyde and stained with corresponding antibodies. **(C)** Cell cycle and morphological analysis. > 200 cells were counted for each time point for each treatment treated as in (B). Average of three independent experiments. Error bars represent standard deviation (SD). **(D)** Representative pictures of control cells and SMC2-depleted cells at different stages of mitosis. DNA (blue), CENP-T (red), α-tubulin (green). Coloured box at the bottom right corners of merge panels correspond to the stages of mitosis in (C). SMC2-depleted mitotic cells have a single chromosome mass adjacent to a malformed mitotic spindle. Stretched chromatin fiber (arrow in i, right). Normal prometaphase-like chromosomes were occasionally observed within the auxin-treated cell population but all these cells were SMC2-mAID-GFP positive. **(E)** Stills from live-cell imaging of SMC2-AID-GFP/CDK1^as^ expressing Pericentrin/AKAP450 centrosomal targeting (PACT)-RFP treated with auxin 3 - 6 h. DNA was stained with SiR-DNA. 3D image stacks were collected at 0.4-μm z increments on a Zeiss Airy microscope every 5 mins. SMC2-AID-GFP/CDK1^as^ cells exited mitosis without apparent chromosome segregation. Chromosomes appeared to be decondensed and centrosome positioned close by in later time points (xii-xiv) Control (no auxin) is shown in Figure 2-figure supplement 1D. Scale bars 5 μm.

The strong chromosome phenotype observed after rapid condensin depletion made it difficult to determine the stage of mitosis at which the cells arrested. We therefore stained a sub-clone of SMC2-AID-GFP cells, expressing CENP-H-GFP from the endogenous allele, with antibodies to histone H3S10ph and INCENP following treatment with vehicle (ethanol) or auxin for 3 h. The signal intensity of H3S10ph is normally high on prometaphase and metaphase chromosomes and progressively decreases after anaphase onset. INCENP is a scaffolding and regulatory subunit of the chromosome passenger complex that localises on chromosome arms and centromeres in early mitosis, then transfers to the central spindle and cleavage furrow during mitotic exit (Carmena et al., 2012).

The H3S10ph level on the SMC2-depleted chromosome mass resembled that on control prometaphase or metaphase chromosomes (Figure 1Ei,iii, 2Aii,iii). Furthermore, INCENP localised on the SMC2-depleted chromosome mass and not on the spindle (Figure 2-figure supplement 1A). These observations suggest that after auxin treatment, SMC2-depleted cells with chromosome clusters were in early mitosis.

These results in SMC2-AID-GFP cells treated with auxin revealed that condensin is required to shape and resolve individual mitotic chromosomes in cultured vertebrate cells. In addition, phenotypic differences between SMC2^OFF^ cells and SMC2-AID-GFP cells depleted of SMC2 supported our hypothesis that the milder defects observed in previously reports were due to the gradual loss of condensin.

### Use of chemical genetics to obtain mitotically synchronised cells lacking condensin

In order to obtain DT40 cultures progressing through mitosis in a synchronous wave, we introduced a CDK1^as^ cDNA into SMC2-AID-GFP cells and then disrupted the endogenous CDK1 alleles using CRISPR/Cas9 technology. 1NMPP1, a bulky ATP analogue, inhibits the mutant CDK1^as^ but no other kinases in these cells. As expected, 1NMPP1 did not inhibit cells from progressing through G_1_, S and G_2_ phases, however, it prevented G_2_ cells from entering mitosis, and forced pre-existing mitotic cells to exit mitosis prematurely (Hochegger et al., 2007). No mitotic cells were detected 30 min after addition of 1NMPP1 (Figure 2-figure supplement 1B). Subsequent washout of 1NMPP1 from the arrested G_2_ cells triggered a rapid and near synchronous wave of mitotic entry and exit (Figure 2C, Figure 2-figure supplement 1C).

Introduction of the CDK1^as^ circuit into SMC2-AID-GFP cells (termed SMC2-AID-GFP/CDK1^as^ cells) allowed us to investigate the effects of depleting condensin either just prior to mitosis or during a mitotic arrest while avoiding confounding effects from previous cell divisions.

A typical experimental time line using SMC2-AID-GFP/CDK1^as^ cells is illustrated in Fiure 2B (see Materials and Methods). All control cells (no auxin) were in interphase at T= 0 min. The mitotic index of the control cells increased gradually from T= 15 min, peaked at T= 60 min, then decreased thereafter (Figure 2C). Prophase cells were observed at T= 15 min but were mostly gone by 30 min. Most mitotic cells at T= 30 - 60 min were in prometaphase or metaphase (Figure 2C). Cells in anaphase and telophase were seen at T= 60 – 120 min. Furthermore, G1 cells were detected 1 h after release from a 1NMPP1 block by DNA content analysis (Figure 2-figure supplement 1C). Thus, mitosis lasts about one hour in control cells after release from a 4 h 1NMPP1 block.

### SMC2-depleted cells accumulate at abnormal prometaphase then exit mitosis without segregating chromosomes

Control cells exhibited prophase chromosome condensation within 15 min of the 1NMPP1 washout (Figure 2C). In contrast, SMC2-depleted cells did not show chromatin condensation until after nuclear envelope breakdown, as previously reported with RNAi or conditional knockouts of condensin (Hudson et al., 2003; Hirota et al., 2004; Ono et al., 2004).

The lack of prophase condensation could explain the slightly lower mitotic index of SMC2-depleted cells at T= 15 min (Figure 2C). However, the mitotic index caught up with that of control cells by T= 30 - 60 min and continued to increase throughout the time course. No recognizable anaphase or telophase was observed in SMC2-depleted cells over the 2 h time course of this experiment confirming the results in asynchronous culture (Figure 2C, D). Nevertheless huge polyploid interphase and mitotic cells were seen following 2 days auxin treatment (Figure 2-figure supplement 1E), suggesting somehow these cells exited the previous mitosis.

Live cell imaging of SMC2-depleted cells showed that these cells exited mitosis without properly segregating chromosomes (Figure 2E) although control cells did (Figure 2-figure supplement 1D). This can explain why we did not observe anaphase or telophase cells following SMC2-depletion.

### SMC2-depleted cells have a malformed mitotic spindle with the chromosome mass to one side

Control chromosomes congress to the middle of a bipolar spindle during late prometaphase and metaphase (Figure 1E, 2Di, 3A, 3B, Figure 3-figure supplement movie 1). In contrast, the single amorphous mass formed by SMC2-depleted chromosomes appeared to be largely excluded from the spindle. A few chromatin fibers protruded from this mass and extended into the spindle with centromeres at their tips (Figure 2Di, 3A, Figure 3-figure supplement movie 2). This suggests that chromosome arms did not follow as their centromeres congressed to the spindle midzone.

**Figure 3.**
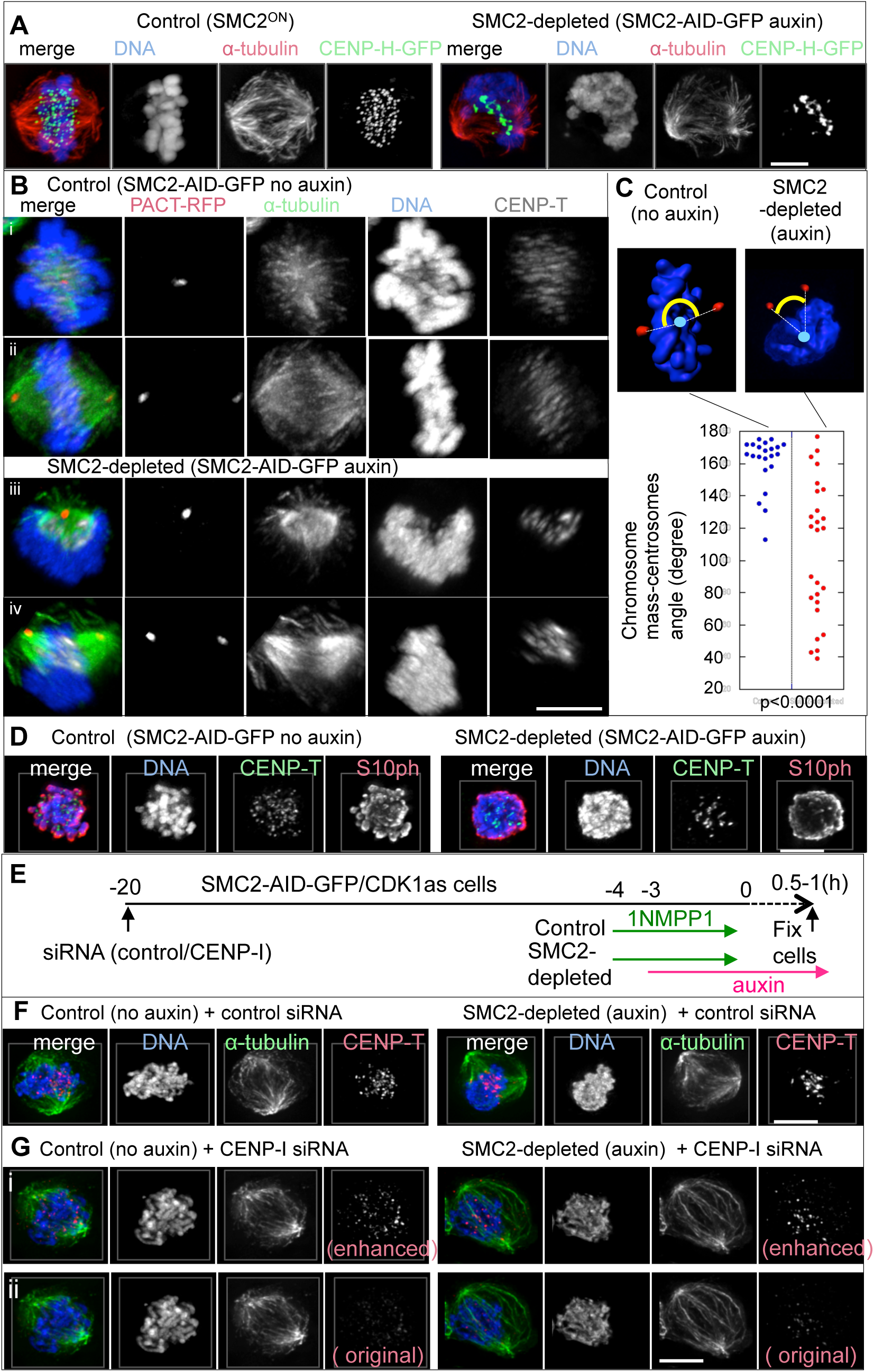
Malformed and mis-positioned chromosomes and mitotic spindle in SMC2-depleted cells. **(A)** SMC2^ON^/CDK1^as^/CENP-H-GFP cells and SMC2-AID-GFP/CDK1^as^/CENP-H-GFP (treated with auxin) were fixed with formaldehyde and stained for α-tubulin (red) and DNA (blue). Images were acquired by structural illumination microscopy. **(B)** SMC2-AID-GFP/CDK1^as^ expressing PACT-RFP was treated with 1NMPP1 for 4 h with or without auxin for 3 h. These cells were fixed with formaldehyde 1 h after 1NMPP1 washout and stained for α-tubulin (green), CENP-T (white), DNA (blue). Cells were 3D-rotated so that two PACT-RFP signals overlap (i, iii) or position in parallel (ii, iv). Note: Signals appeared to be fuzzier and stretched due to 3D-rotation. **(C)** SMC2-AID-GFP/CDK1^as^ expressing PACT-RFP was treated as (B) and stained for DNA. Angles of chromosomes and centrosomes were calculated as described in Materials and Methods. Control (n= 22), SMC2-depleted (n= 26). P value was <0.0001 (two-tailed Mann-Whitney U Test). **(D)** Control and auxin-treated cells treated with nocodazole for 1 h and fixed with formaldehyde. DNA (blue), CENP-T (green), H3S10ph (red). **(E)** Experimental timeline for CENP-I depletion. **(F, G)** SMC2-AID-GFP/CDK1^as^ cells were transfected either with control siRNA oligo **(F)** or CENP-I oligo **(G).** Cells were further treated with ethanol or auxin. The chromosome mass became symmetrical in SMC2-depleted cells when kinetochore binding to microtubules was perturbed. CENP-T signals were enhanced in (i) in order to visualise the position of kinetochores. Non-enhanced images are also shown (ii). DNA (blue), CENP-T (red), α-tubulin (green) Scale bars 5μm.

The mitotic spindle in SMC2-depleted cells was bipolar but often twisted or asymmetric (Figure 1E, 2D, 3A, 3B, Figure 3-figure supplement movie 2). Mitotic spindles and chromosomes in SMC2-depleted cells did not position properly between the two centrosomes, which were visualised using PACT-RFP (<u>p</u>ericentrin/<u>A</u>KAP450 <u>c</u>entrosomal <u>t</u>argeting - Figure 3A, B). In order to quantify the relative positions of chromosomes relative to the mitotic spindles, angles between chromosomes and centrosomes in late prometaphase or metaphase cells were measured using SMC2-AID-GFP/CDK1^as^ cells expressing PACT-RFP (see Materials and Methods). The angle between the centre of mass of the chromosomes and the two centrosomes was between 155-180 degrees in most control cells (18 out of 22) (Figure 3B,C). In contrast, this angle was less than 90 degrees in nearly half of the SMC2-depleted cells (13 out of 27).

Forces exerted by spindle attachments exacerbated the structural defects of mitotic chromosomes and altered the relative position and shape of the chromosomes and spindle. When microtubules were depolymerized by nocodazole treatment in control cells, rod-shaped individual mitotic chromosomes could be recognized (Figure 3D). In contrast, individual chromosomes could not be resolved in SMC2-depleted cells treated with nocodazole although the chromosome mass became more symmetrical and protruding chromatin fibers disappeared.

CENP-I is a kinetochore protein whose depletion by RNAi causes defects in kinetochore assembly, visualized as decreased CENP-T levels at kinetochores (Figure 3E-G). Microtubules do not bind tightly to such compromised kinetochores (Nishihashi et al., 2002; Hori et al., 2008a). In CENP-I-depleted cells, the morphologies of SMC2-depleted chromosomes were still aberrant, however the chromosome mass was symmetrical and overlapped the mitotic spindle, which also adopted a symmetrical shape (Figure 3G).

These experiments confirm that condensin is required to shape mitotic chromosomes and reveal that mitotic spindle forces further deform the amorphous SMC2-depleted chromosomes, presumably because SMC2-depleted chromosomes lack structural integrity in arms and in pericentromere regions as reported previously (Oliveira et al., 2005; Gerlich et al., 2006; Bouck and Bloom, 2007; Ribeiro et al., 2009). We also confirmed that lack of structural integrity in SMC2-depleted chromosomes results in mispositioning of the chromosome mass and spindle malformations (Wignall SM 2003, Ono T 2004).

### SMC2-depleted kinetochores fail to make stable attachments to microtubules

The CENP-H-GFP signals on SMC2-depleted kinetochores that stretched out from the chromosome mass were round (21 out of 21 centromeres) and resembled those in control metaphase cells, where correlative light and electron microscopy (CLEM) revealed a typical trilaminar kinetochore structure with attached microtubules (Figure 4A arrows, B). In contrast, the CENP-H-GFP signal on SMC2-depleted kinetochores inside or at the edge of the chromosome mass was often deformed (Figure 4C yellow arrowheads, see also Figure 2A and Figure 3-figure supplement movie 2). Indeed the trilaminar structure was less evident following SMC2 depletion (Figure 4A, B). The shape of kinetochores may be distorted due to lack of structural integrity of the underlying chromatin as previously suggested (Samoshkin 2009).

**Figure 4.**
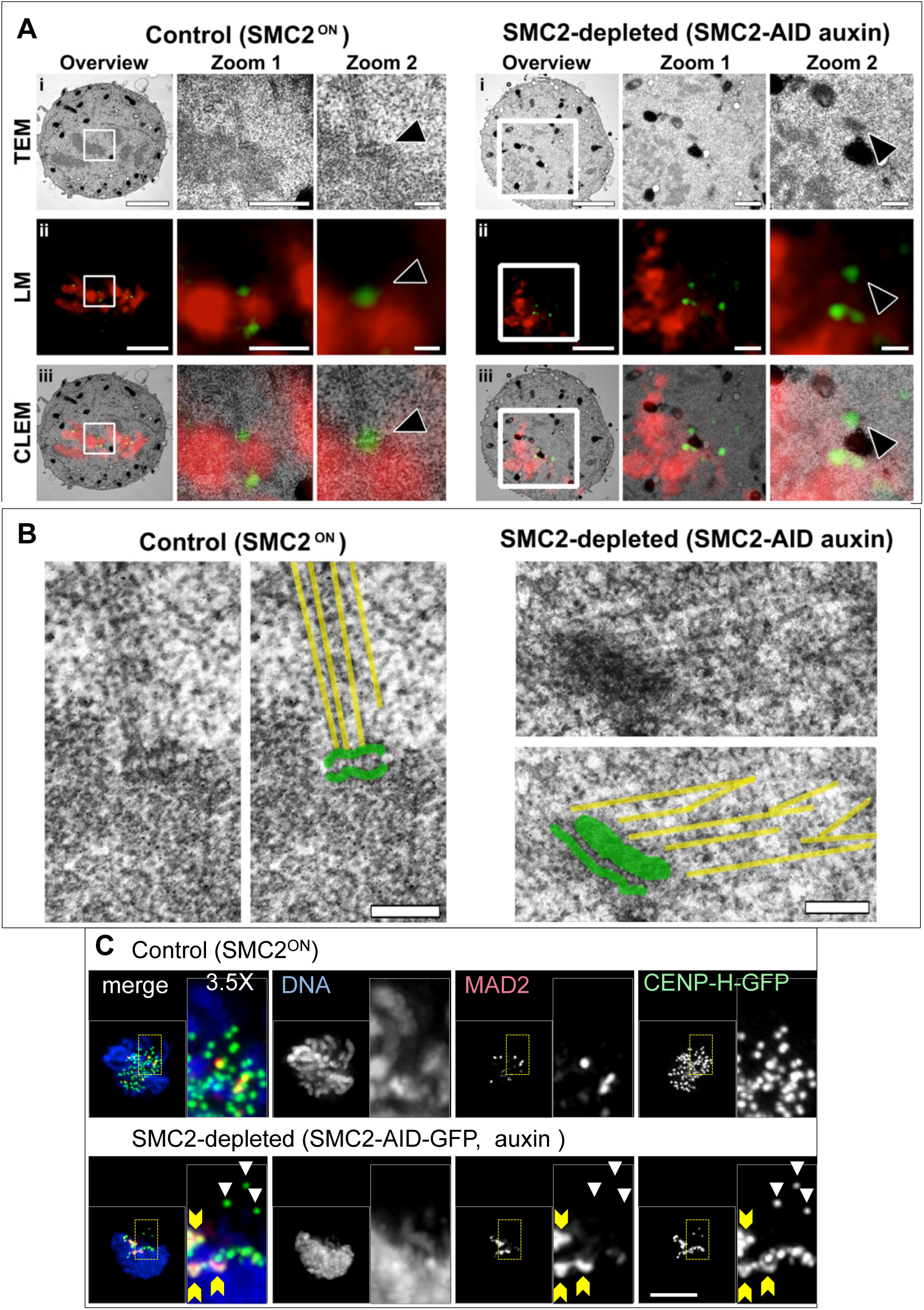
Compromised kinetochore-microtubule attachments in SMC2-depleted cells. **(A)** Correlative light and transmission electron microscopy of control and SMC2 depleted cells. Images show three progressive zooms of the white-boxed region, for TEM (i), light microscopy of DAPI (red) and CENP-H-GFP (green) (ii) and a correlative overlay of both the physical and optical sections (iii). Arrow in zoom 2 points to a clearly defined trilaminar structure. Scale bars A - 3μm, 1μm, 200nm for overview, zoom 1 and zoom 2 respectively. **(B)** Enlargement of kinetochores shown in Figure 4. Microtubules (yellow lines) and kinetochore (green) were overlaid on the TEM pictures, and show microtubule attachment. Scale bar 200 nm. **(C)** SMC2^ON^/CDK1^as^/CENP-H-GFP cells and SMC2-AID-GFP/CDK1^as^/CENP-H-GFP (treated with auxin) were treated with 1NMPP1 for 4 h and fixed with formaldehyde 1 h after 1NMPP1 washout. Strong MAD2 staining on the centromeres inside of the SMC2-depleted chromosome mass (yellow arrowheads) is not observed on centromeres outside of the chromosome mass (white arrows). DNA (blue), MAD2 (red), CENP-H-GFP (green). Scale bar 5 μm.

Strong MAD2 signals were seen in almost all SMC2-depleted late prometaphase cells (24 out of 25 cells at 1 h after release from 4 h 1NMPP1 block) (Figure 4C). In contrast, approximately half of control late prometaphase cells showed strong MAD2 signals (13 out of 25 cells) and other cells showed only weak MAD2 signal (11 out of 25 cells). This suggested that these MAD2 positive centromeres lack stable bipolar attachments and that the prometaphase accumulation of SMC2-depleted cells (Figure 2C) might be due to an unsatisfied spindle assembly checkpoint (SAC).

Indeed, silencing of the SAC by Mad2 RNAi caused both SMC2-depleted and control cells to enter anaphase with unaligned chromosomes (Figure 4-figure supplement 1). Mitotic exit was confirmed by INCENP staining on the central spindle and by reduced Cyclin B2 staining (Figure 4-figure supplement 1A, Bi,iii,iv). Approximately 80% of SMC2-depleted cells treated with MAD2 siRNA showed highly uneven chromosome segregation with chromosome bridges and centromeres trapped in the intercellular bridge (Figure 4-figure supplement 1A, B, D). Live cell imaging showed that all SMC2-depleted daughter cells treated with MAD2 siRNA fused after an abortive cytokinesis (Figure 4-figure supplement 1E).

Thus, SMC2-depleted kinetochores can attach to microtubules but are defective in forming stable amphitelic attachments.

### SMC2 is required for intrinsic chromosome organization but not for mitotic chromatin compaction

To better understand the consequences of rapid SMC2 depletion on chromosome morphology we performed an ultrastructural analysis using 3D-CLEM with SBF-SEM (serial block face scanning electron microscopy) and digital reconstructions (Booth et al., 2016).

To establish a default set of control parameters, the parental wild type DT40 cell line was first analysed. The entire chromosome complement of a metaphase control cell, was reconstructed from 140 serial sections and modeled using Amira (Figure 5A). The presence of two pairs of centrioles surrounded by PCM and flanking the aligned chromosomes confirmed that the cell was indeed in metaphase. Using separation algorithms we identified 74 discrete chromosome units (Figure 5Av-vi), with a combined volume of 75 μm^3^ and surface area of 424 μm^2^ (Figure 5D). The normal DT40 karyotype has 80 chromosomes, though scoring of the smallest microchromosomes can be difficult (Sonoda et al., 1998; Molnar et al., 2014).

**Figure 5.**
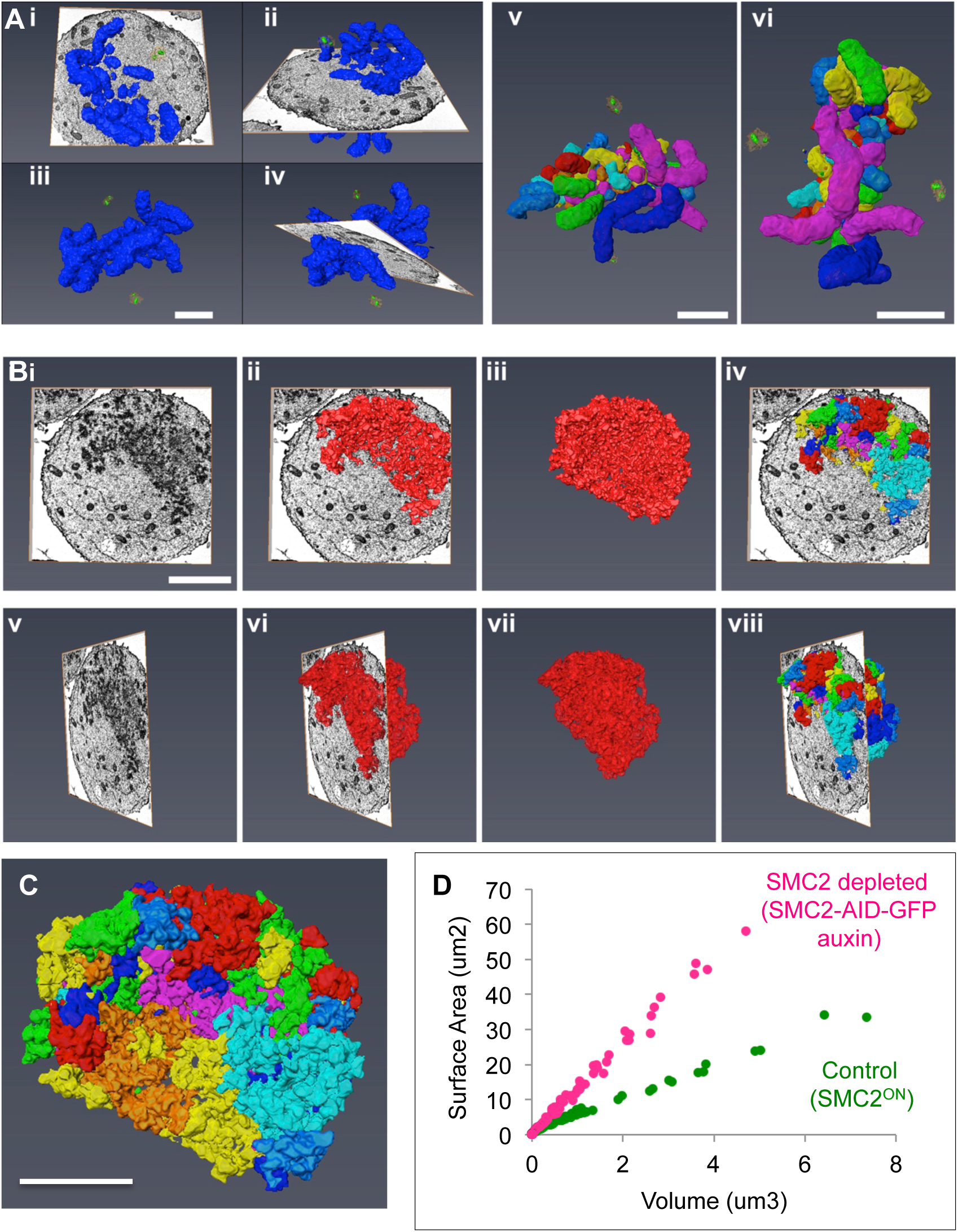
3D-EM shows that condensin is required for chromosome organization, but not for chromatin compaction. WT and condensin depleted cells were prepared for 3D SBF-SEM analysis. **(A)** A control cell with all chromosomes modeled (indigo) is shown at different angles, with and without the EM orthoslice (i – iv). Centrioles (green) and Pericentriolar material (orange) are also shown. Chromosomes were separated into individual units (multi-coloured) (v-vi). **(B)** A condensin depleted cell with all chromosomes modeled (red), shown at different angles, with and without the EM orthoslice (i–iii, v-vii). Separated chromosomes traversing an EM orthoslice (iv and viii). **(C)** SMC2-depleted chromosomes were separated into discrete units. **(D)** 2D scatter plot of surface area versus volume for individual chromosomes in A (control, green) and B (condensin depleted, magenta). Scale bar 5 μm.

A similar analysis was next performed on a cell following rapid SMC2 depletion (Figure 5B). Remarkably, despite the clear disorganization of the chromatin mass, it was possible to resolve 84 chromosome units (Figure 5C), with a combined volume of 71 μm^3^ and surface area of 931 μm^2^ (Figure 5D). Interestingly, although both the chromosome number and combined volumes were similar (74 vs 84 and 75 μm^3^ vs 71 μm^3^), the surface area had more than doubled following SMC2 depletion (424 μm^2^ vs 931 μm^2^). This suggests that chromatin compaction measured as total chromatin volume was near normal, but that the shape of the chromosomes as revealed by the surface area was severely compromised. By ranking the chromosomes according to their volumes, we could confirm that individual chromosomes from the depleted cell had a similar volume, but a far greater surface area than wild-type chromosomes (Figure 5D).

This analysis confirms that, as previously suggested (Vagnarelli et al., 2006), condensin has a key role in establishing a normal mitotic chromosome architecture, but is not essential for volume-wise reduction of chromatin (chromatin compaction) from interphase to mitosis.

### Condensin is essential for normal intrinsic architecture of condensed mitotic chromosomes

Use of the intrinsic mitotic chromosome structure (IMCS) assay previously established that condensin is required to establish a normal mitotic chromosome architecture (Hudson et al., 2003) (Figure 6A). In this assay, cells were treated with nocodazole for 12 h before exposure to a hypotonic buffer (Figure 6-figure supplement 1A). To test condensin’s role in establishing or maintaining the structural memory of mitotic chromosomes, respectively, cells were treated with auxin either for the entire 12 h (SMC2 depleted before mitotic chromosome assembly) or for the last 2 h (SMC2 depleted after mitotic chromosome assembly). Auxin was omitted from controls so that SMC2-mAID-GFP was continuously present.

**Figure 6.**
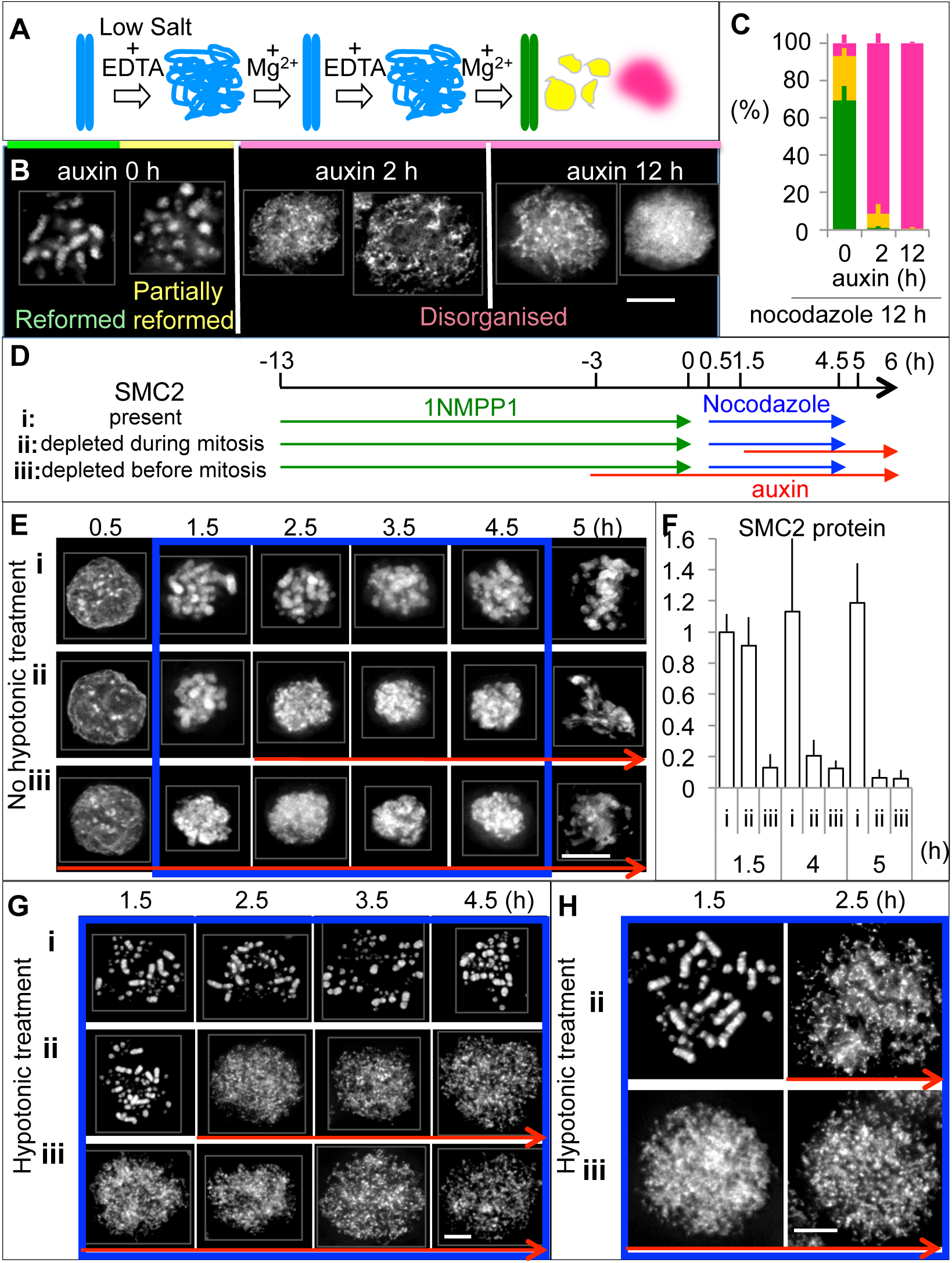
SMC2 is required to maintain the structure of mitotic chromosomes. **(A)** The intrinsic mitotic chromosome structure (IMCS) assay. Cells were blocked in mitosis with nocodazole. Removal of cations in hypotonic buffer induced the unfolding of chromatin to the level of 10 nm fibres. Addition of Mg^2+^ triggered the shrinking and refolding of chromatin. The shape of chromosomes after two cycles of unfolding and folding was classified as reformed (green), partially reformed (yellow) or disorganised (magenta). **(B)** Representative images of DAPI-stained mitotic chromosomes at the end of the assay. Scale bar 5 μm. **(C)** Quantification of chromosomes at the end of the assay. The graph shows the mean and SD of three independent experiments with > 50 cells/each treatment counted. **(D)** Experimental procedure for determining the role of condensin in establishing and maintaining mitotic chromosome structure. Cultures were treated with 1NMPP1 for 13 h. After 1NMPP1 washout, cultures were treated with nocodazole for 4 h to hold the cells in mitosis (T= 0.5 – 4.5 h). The mitotic index reached approximately 80 % after 1 h of nocodazole addition and did not change until nocodazole washout at 4.5 h. Cells were fixed with formaldehyde at indicated time points and stained with DAPI before > 100 cells/each treatment’s time point were counted. Mean and SD of three independent experiments are shown at F and Figure 6-figure supplement 2B-D. **(E)** Cells were fixed with formaldehyde/PBS without hypotonic treatment at the indicated time points and stained with DAPI. **(F)** The amount of SMC2 remaining was measured by immunoblot analysis and normalized to the level of α-tubulin. **(G, H)** Cells were treated with 75 mM KCI at 37 °C for 5 min prior to ice-cold methanol/acetic acid fixation and DAPI staining.

For the IMCS assay, chromosomes were subjected to two cycles of unfolding and refolding, then their shape was scored in one of three categories: reformed, partially reformed or disorganised (Figure 6A, B). Approximately 90 % of control cells (minus auxin) showed either reformed or partially reformed chromosomes (Figure 6B, C). In contrast, cells depleted of SMC2 prior to chromosome assembly (auxin for 12 h) exhibited only disorganized chromatin after this treatment. Remarkably, a similar result was obtained for > 90 % of cells depleted of SMC2 only after mitotic chromosome formation (auxin for 2 h). Depletion of SMC2-mAID-GFP after auxin addition was confirmed by immunoblot analysis (Figure 6-figure supplement 1B, C). Thus, the structural memory of mitotic chromosomes is not maintained in the absence of condensin, even when SMC2 is degraded *after* mitotic chromosomes have formed.

We next exploited the ability of SMC2-AID-GFP/CDK1^as^ cells to enter mitosis in a synchronous wave in order to follow the process of chromosome formation in the presence or absence of condensin. We also characterized chromosomes depleted of SMC2 during mitosis (Figure 6D, Figure 6-figure supplement 2A).

All cells in the culture were in interphase at the end of a 13 h 1NMPP1 treatment (T≈ 0.5 h) (Figure 6E). Approximately 80% of those cells entered mitosis within 1h after 1NMPP1 washout (T= 1.5 h). The mitotic index then remained almost unchanged until nocodazole washout (T= 4.5 h).

Mitotic chromosomes were readily visible in control cells during nocodazole treatment (Figure 6Ei, 6Gi: T= 1.5 - 4.5 h). Chromosomes began to align at a metaphase plate by 30 minutes after nocodazole washout at T = 4.5 h (Figure 6Ei). Most of these cells exit mitosis at T= ≤6 h, as revealed by a reduction in Cyclin B2 levels (Figure 6-figure supplement 2C, Ei, Fi).

Depletion of SMC2 after mitotic chromosome formation revealed that condensin is required to maintain chromosome morphology in mitosis. Rod-shaped chromosomes were clearly visible prior to auxin addition at T= 1.5 h, but became much less distinct at T= 2.5 h, within 1 h of auxin addition (Figure 6Eii). After hypotonic treatment, chromosomes appeared normal at T= 1.5 h, but were completely disorganised within 1 h of auxin addition at T= 2.5 h (Figure 6Gii, 6Hii).

For cells in which SMC2 had been depleted prior to mitotic entry, mitotic chromosome structure was defective at all time points following release from the 1NMPP1 block (T= 1.5 – 4. 5 h - Figure 6Eiii). Without hypotonic treatment, these SMC2-depleted mitotic chromosomes appeared as a condensed ball. In cells treated with hypotonic buffer prior to fixation, SMC2-depleted mitotic chromosomes formed a disorganized mass of dispersed chromatin at all time points (Figure 6Giii, 6Hiii).

The morphology of SMC2-depleted mitotic chromosomes and subsequent chromosome mis-segregation defects were indistinguishable regardless of whether SMC2 was depleted prior to mitotic entry or during mitosis after chromosomes had formed. This confirms for somatic cells the early observation (Hirano and Mitchison, 1994) and recent studies with mouse oocytes and Drosophila embryo (Houlard 2015, Piskadlo 2017), that condensin is necessary to maintain the structure of mitotic chromosomes throughout mitosis.

### Titration of condensin levels reveals differential requirements for SMC2 in mitotic chromosome formation and chromosome segregation

Condensin is required both for maintenance of chromosome architecture and for sister chromatid segregation at anaphase. To further explore the functional link between these two processes, we used auxin treatments to produce cultures with a graded series of condensin concentrations. This was possible because the abundance of an AID tagged protein can be manipulated by varying the auxin concentration (Nishimura et al., 2009).

Auxin concentrations yielding a range of partial depletions of SMC2-mAID-GFP were selected based on data obtained from flow cytometry, microscopy, and immunoblotting analysis (Figure 7A, B). SMC2-AID-GFP/CDK1^as^ cells were treated with 1NMPP1 for 4 h and with various concentrations of auxin for 3 h. After release from 1NMPP1 to allow entry into mitosis, cells were fixed every 30 min and analysed as in Figure 2. Auxin was omitted from wash and release media apart from 125/250 μM auxin treated cells because we noticed that in mitotic cells even low auxin concentrations eventually resulted in depletion levels equivalent to those seen with 125/250 μM auxin treatment. This is likely due to the reduction of protein synthesis during mitosis, such that even a low degradation activity could eventually deplete the AID-tagged protein.

**Figure 7.**
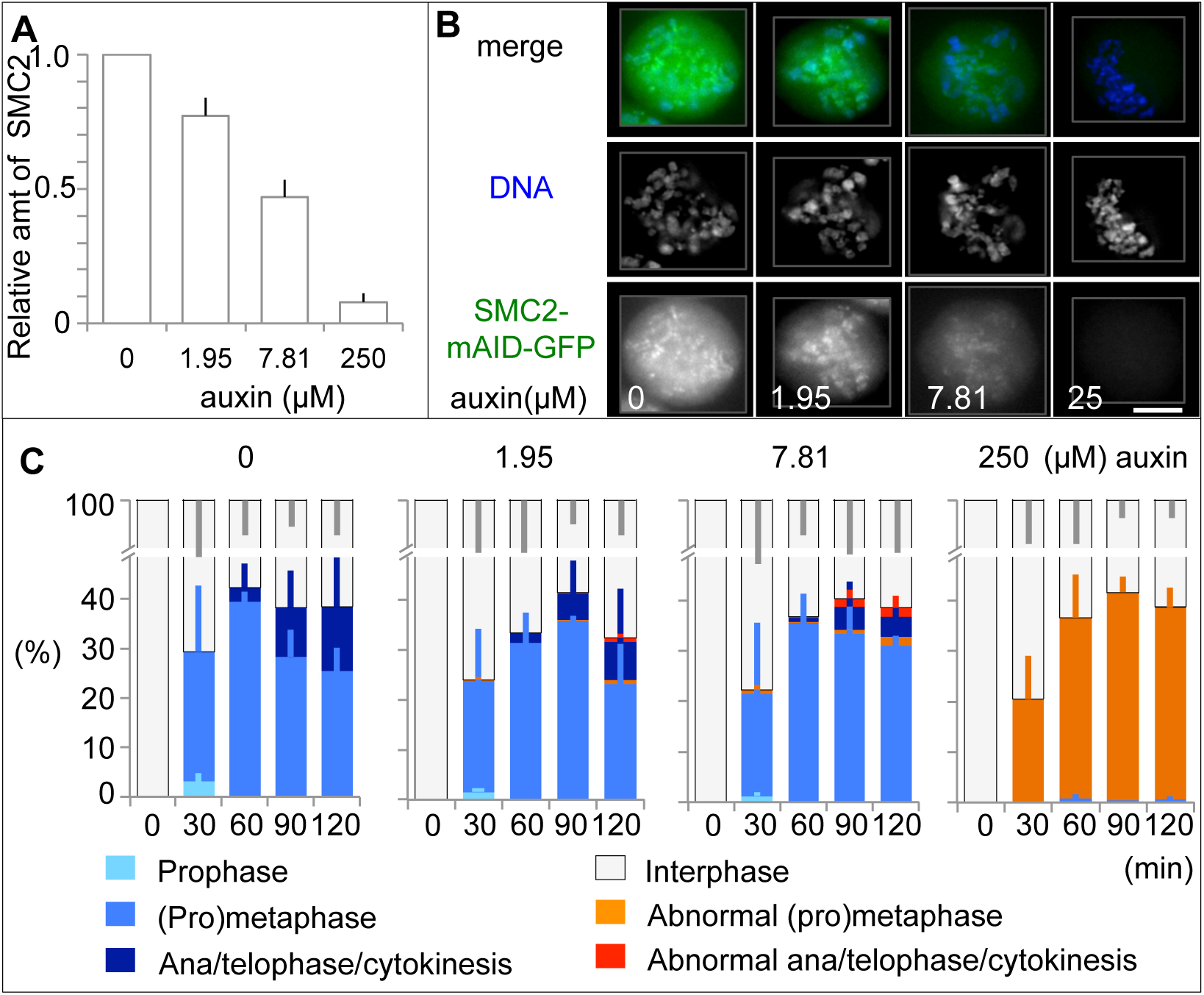
Cells partially depleted of SMC2 show chromosome mis-segregatlon. **(A)** The amount of SMC2 remaining in asynchronous culture upon 4 h auxin treatment was measured by immunoblot analysis and normalized to the level of α-tubulin. Mean and SD of three independent experiments are shown. **(B)** Representative images of SMC2-AID-GFP cells at prometaphase upon 4 h auxin treatment with various concentrations of auxin. SMC2-mAID-GFP (green). Scale bar 5 μm. **(C)** Effect of various auxin concentrations on mitotic progression of SMC2-AID-GFP cells. Experimental procedure as in Figure 2B. Cells were fixed every 30 min after 1NMPP1 washout. Auxin was omitted from the media used for the wash and release except in the case of cells treated with 250 μM auxin. Mean and SD of three independent experiments. > 200 cells were counted for each time point of individual treatments.

Immunoblot analysis showed that cellular levels of SMC2 fell to ^~^75% or ^~^45% of those in wild-type cells after addition of 1.95 μM or 7.81 μM of auxin, respectively (Figure 7A). (The absolute levels vary between experiments but the relative levels are preserved.) Reduction of SMC2 by 25% had little impact on either the morphology or segregation of mitotic chromosomes (Figure 7C). However, chromosome segregation defects were observed when SMC2 fell to ^~^55 % of wild type levels. In those cells, prometaphase chromosomes were slightly wider but individual rod-shaped chromosomes were still discernible. Despite this near normal morphology, some chromatin bridges were observed when the cells entered anaphase and telophase.

We next asked whether the observed defects in sister chromatid segregation were due to defects in the intrinsic structure of the chromosomes or to another function of condensin. We exposed SMC2-AID-GFP/CDK1^as^ cells to 1NMPP1 for 4 h plus various concentrations of auxin for 3 h, then washed out the 1NMPP1 to allow cells to enter mitosis (Figure 8A). At 45 min after 1NMPP1 washout (when cells were in prometaphase), cells were subjected to the IMCS assay as in Figure 6A. Later, at 90 min after IN MPP1 washout (when cells were in anaphase or telophase), cells were fixed and stained to examine sister chromatid segregation. For cells treated with 125 μM auxin we did not observe typical anaphase or telophase cells, so we instead scored chromatin bridges at cytokinesis.

**Figure 8.**
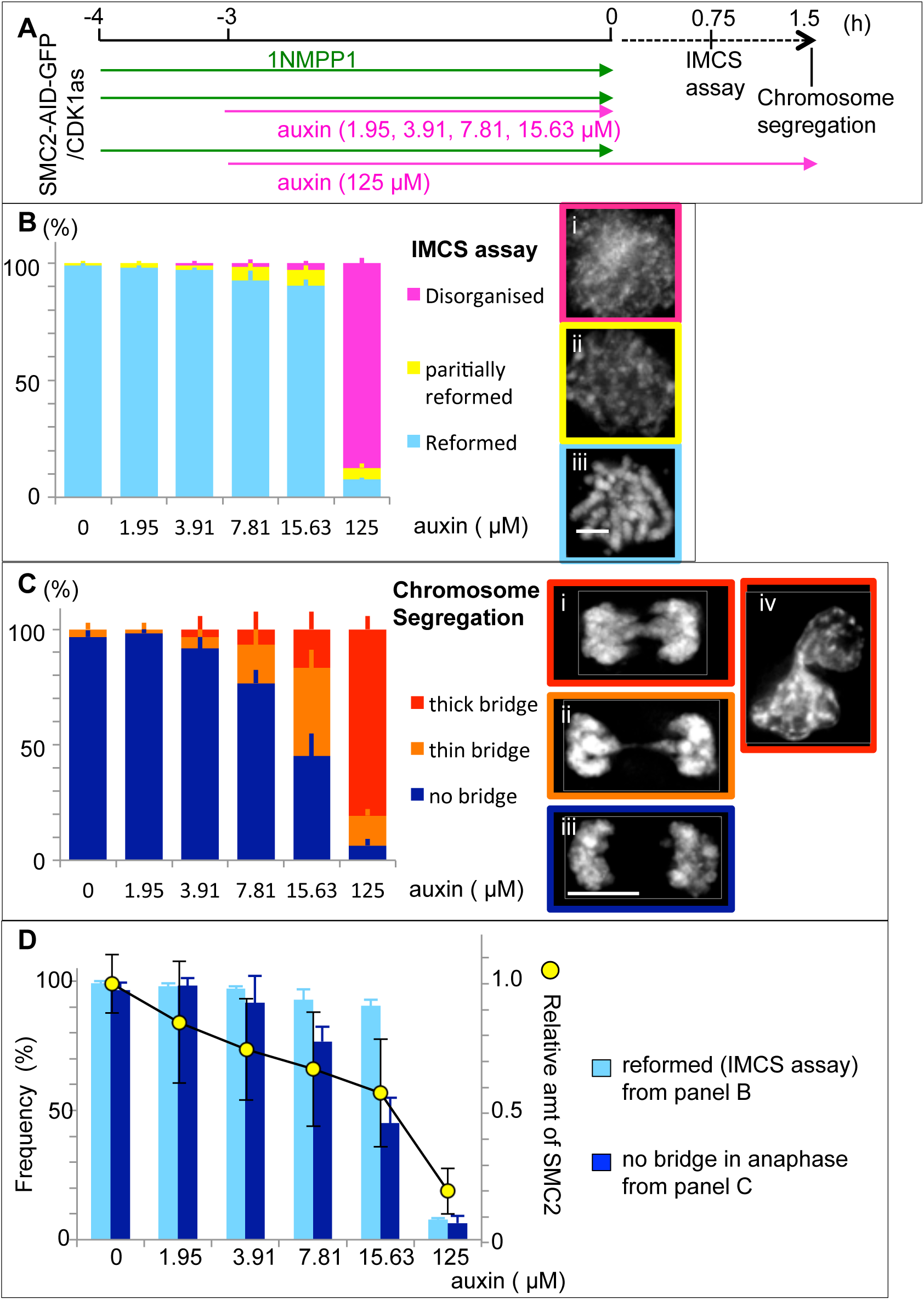
Differing amounts of condensin are required for chromosome assembly and for segregation. **(A)** Experimental protocol. **(B)** 45 min after 1NMPP1 washout, cells were subjected to the IMCS assay. Representative images are shown. > 100 cells/each treatment were counted. Mean and SD of three independent experiments. **(C)** 90 min after 1NMPP1 washout, cells were fixed and stained with anti α-tubulin antibody and DAPI. Representative images are shown. > 20 anaphase or telophase cells/each treatment were counted. Cultures treated with 125 μM auxin did not show anaphase/telophase cells, so cells in early cytokinesis (iv) were analysed. Mean and SD of three independent experiments. **(D)** Overlay of key phenotypes from the experiments of (B, C) plotted against the residual levels of SMC2-mAID-GFP measured by immunoblotting and normalized to the level of α-tubulin in cells treated with the indicated concentration of auxin.

This experiment revealed very clear differences in the threshold requirements for condensin in chromosome architecture and sister chromatid segregation. In the IMCS assay, mitotic chromosome structures reformed in 90% of cells when SMC2 protein levels had fallen to 60% of wild type (15.63 μM auxin – Figure 8B, D). In contrast, only 45% of cells successfully segregated chromosomes without chromosome bridges under these conditions (Figure 8C, D).

Collectively, these results reveal that higher levels of condensin are required for sister chromatid resolution and anaphase chromosome segregation than for formation of a normal mitotic chromosome architecture, as assayed by the IMCS assay.

### Partial depletion of condensin disrupts the chromosome axes

Use of the auxin titration series revealed that even under conditions of partial condensin depletion where the IMCS assay reports that chromosome architecture is near normal, the organization of the chromosome axes is strongly affected. When condensin levels have been depleted by as little as 50%, the distribution of condensin and topo IIα is disrupted and the proteins appear to diffuse throughout the chromosomes. This cannot be observed in 125 μM auxin, where condensin is effectively gone and topoisomerase IIα is no longer visible on the chromatid (Figure 9A). Diffuse chromosomal localisation of topoisomerase IIα has been reported previously following conventional condensin depletion (Hudson 2003, Coelho 2003). The anaphase bridges observed under partial depletion of SMC2 could be explained by incomplete decatination and/or possibly recatination of sister chromatids due to disruption of chromosome axis and diffuse localisation of topoisomerase IIα as recently reported (Piskadlo 2017). Furthermore, it could explain why the chromosome segregation defects have been observed as the most prominent and universal defect when condensin depletion occurred with slower kinetics.

**Figure 9.**
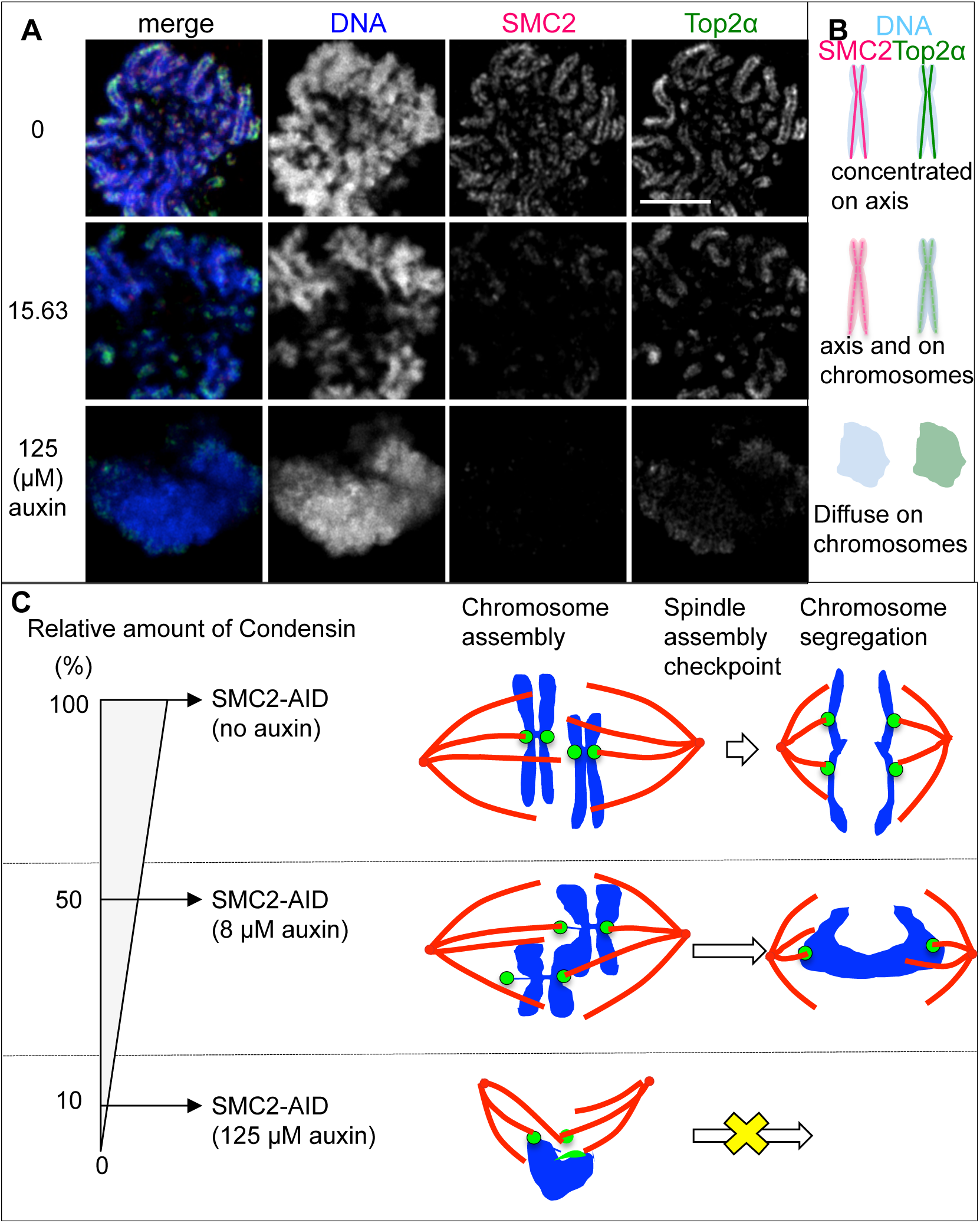
Effects of lowering condensin levels on chromosome structure and segregation. **(A)** SMC2-AID-GFP/CDK1^as^ cells were treated with 1NMPP1 for 13 h and with the indicated amounts of auxin for the final 3 h. Auxin was omitted from media used for the wash and release except for cells treated with 125 μM auxin. 30 min after 1NMPP1 washout, cells were collected, rinsed with PBS and fixed with cold methanol/acetic acid. DNA (blue), SMC2 (red), and Top2α (green). Scale bar 5μm. **(B)** Diagram summarizing the changes in protein distribution seen in (A). **(C)** Summary of the effects of differing condensin levels on mitotic DT40 cells. DNA (blue), centromeres (green), mitotic spindle (red).

## DISCUSSION

Mitotic chromosome condensation is a complex process, involving removal of interphase chromatin structures (TADs and compartments), two-to-three-fold compaction of the chromatin, formation of rod-shaped structures and sister chromatid resolution (Vagnarelli, 2012; Naumova et al., 2013). The role(s) of condensin in these processes remain unclear. Here we show that condensin is not required for mitotic chromatin volume compaction and that different threshold levels of condensin are required for establishment of mitotic chromosome architecture and for chromatid segregation in anaphase.

There has been a long-standing controversy as to what are the primary mitotic defects after condensin depletion. Prior to the discovery of condensin, it was reported that SMC proteins were needed to both assemble and maintain mitotic chromosomes in Xenopus egg extracts (Hirano and Mitchison, 1994). A recent study of mouse meiotic cells and *Drosophila* embryos reached a similar conclusion (Houlard et al., 2015), but in between, many studies of somatic cells reported that rod-shaped chromosomes could form even when condensin had been significantly depleted (Bhat et al., 1996; Steffensen et al., 2001; Hagstrom et al., 2002; Somma et al., 2003; Hudson et al., 2003; Coelho et al., 2003; Vagnarelli et al., 2006; Maddox et al., 2006; Samoshkin et al., 2009). Interestingly, those studies agreed that although chromosomes could form in cells depleted of condensin subunits, anaphase sister chromatid separation was seriously impacted and prominent chromatin bridges were commonly seen.

The present study combined use of auxin inducible degron (AID) technology for rapid depletion of condensin with CDK1^as^ alleles to obtain improved synchrony of mitotic entry. SMC2-AID-GFP/CDK1^as^ cells showed more homogenous and more severe defects than those seen in SMC2^OFF^ conditional knockout cells. Interestingly, residual levels of SMC2 were similar in both systems (5-7 % of wild type cells). Importantly, the AID system appears to function particularly well in somatic mitotic cells, where protein translation is reduced. Thus, the residual levels of condensin in mitotic cells may be lower compared to those in G2 phase.

We hypothesize that previous discrepancies in chromosome morphology observed with differing experimental protocols did not simply reflect the extent of condensin depletion, as previously speculated, but instead the time taken to deplete condensin, and the time allowed for cells to up-regulate compensatory pathways.

Use of SMC2-AID-GFP/CDK1^as^ cells allowed us to obtain >10^8^ cells with ≥ 80% purity in G2, prophase and prometaphase without microtubule inhibitors. We could distinguish the roles of condensin at mitotic entry, during metaphase and during mitotic exit because SMC2 could be depleted just prior to mitotic entry and also within mitosis in a time restricted manner. We found that mitotic chromosomes depleted of condensin after their assembly were morphologically and structurally indistinguishable from chromosomes assembled in the absence of condensin. This is consistent with observations in the Xenopus egg extract system (Hirano and Mitchison, 1994). It is tempting to speculate that one essential role of condensin is to lock-in the architecture of mitotic chromosomes and prevent other factors such as topoisomerase IIα from promoting aggregation or random (de)catenation of the chromatin.

Two recent studies showed that meiotic chromosomes in mouse oocytes or mitotic chromosomes in early *Drosophilia* embyos rapidly depleted of condensin lost structural integrity and were morphologically aberrant (Houlard et al., 2015) (Piskadlo 2017). Importantly in both of these studies and in the present work, the resulting chromatin did not look like interphase, but was highly compacted although the usual rod-like morphology was lost. The three studies thus clearly show that condensin is essential for shaping and maintaining individual mitotic chromosomes *in vivo* in embryos and in somatic cells. These results are consistent with the original studies with *Xenopus* egg extract systems and with studies in *S. pombe* (Saka et al., 1994). Thus, this role of condensin is evolutionally conserved.

In cells rapidly depleted of SMC2, individual chromosomes could not be distinguished by conventional fluorescence microscopy within the single mass of mitotic chromatin. Indeed, the first impression from conventional electron microscopy was of a shattered mass of condensed chromatin. However, ultra-structural analysis using 3D-CLEM with SBF-SEM and digital reconstructions (Booth et al., 2016) successfully resolved the chromosome mass into units that apparently correspond to individual chromosomes. Moreover, the total volume of SMC2-depleted chromosomes was similar to that of wild type chromosomes at metaphase even though the surface area was >4-fold greater.

This result confirmed our previous hypothesis that condensin is dispensable for mitotic chromatin volume-wise compaction, but required for structural organization of the condensed chromosomes (Vagnarelli et al., 2006; Samejima et al., 2012). Surprisingly, mitotic chromosomes can still form after conventional depletion of all the three major components of the mitotic chromosome scaffold (condensin, topoisomerase IIα and KIF4A) (Samejima et al., 2012). Furthermore, extensive proteomic studies failed to identify any other major structural components of mitotic chromosomes with a likely role in promoting mitotic chromatin compaction (Ohta et al., 2010; Samejima et al., 2015). These results and other studies suggest that chromatin compaction in mitosis is mediated by factors intrinsic to the chromatin, possibly including histone post-translational modifications (Markaki et al., 2009; Georgatos et al., 2009) (Zhiteneva et al, 2017).

In addition to the obvious effects on chromosome architecture, rapid depletion of SMC2 led to spindle defects that included abnormal exclusion of chromosomes from the mitotic spindle as well as the shape of the mitotic spindle itself. One characteristic of condensin depletion is the presence of thin chromatin fibers as centromeres stretch out from the bulk of the chromatin. Condensin is essential for establishing the proper elasticity of centromeric heterochromatin, an important determinant of kinetochore movements under tension (Ribeiro et al., 2009; Samoshkin et al., 2009; Jaqaman et al., 2010). The stretched fibers correspond to centromeric heterochromatin that becomes abnormally compliant after condensin depletion. This heterochromatin is stretched by spindle forces acting on kinetochores. Interfering with microtubule attachments at kinetochores restores, to some extent, the chromosome morphology.

Regarding the shape of the spindle, it has long been known that tension stabilises kinetochore-microtubule attachments (Nicklas and Koch, 1969; Akiyoshi et al., 2010) (Miller MP et al, 2016). We postulate that either the lack of tension, compromised kinetochore structure, or possibly mis-localised aurora B kinase activity due to the loss of condensin affects kinetochore microtubule attachments. Furthermore, lack of structural integrity in chromosomes appeared to cause their incorrect positioning relative to centrosomes and the mitotic spindle. It is tempting to speculate that the centromeres are under force from both the mitotic spindle and the bulky chromosome mass. This may in turn produce the abnormal mitotic spindle morphology observed in SMC2-depleted cells and possibly promote incorrect kinetochore-microtubule attachments (Saunders and Hoyt, 1992) (Wignall SM 2003, Ono T 2004).

Availability of an AID degron allele allowed us to examine the effects of creating a hypomorphic series in which levels of SMC2 were lowered to differing extents. The residual amounts of AID tagged proteins are inversely correlated with the auxin concentration (Nishimura et al., 2009). Use of a titration series allowed us to demonstrate that higher levels of condensin are required for successful anaphase sister chromatid separation than for establishing the metaphase chromosome architecture as defined by our IMCS assay. In other experiments, cells rapidly depleted of either condensin I or II complex alone still form morphologically distinct individual mitotic chromosomes but exhibit chromosome segregation defects, indicating the requirement for a higher amount of condensin at anaphase (data not shown). This could explain why severe mitotic chromosome assembly defects were not reported in the experimental systems with gradual depletion of condensin. Failures in cytokinesis during previous cell divisions could have interfered with the observation of morphological defects at the next mitosis.

Why is the threshold amount of SMC2 higher for segregating chromosomes? Partially lowering condensin levels appears to disrupt the orderly distribution of condensin and topo IIα along the chromatid axes. This altered distribution of the two proteins may either cause anaphase chromosomes to become mechanically more fragile or interfere with the decatenation of sister chromatids. The existence of different threshold levels of condensin is consistent with the idea that condensin’s roles in chromosome condensation and sister-chromatid separation are different (Sullivan et al., 2004; D’Amours et al., 2004).

In summary, we show here that the architecture of mitotic chromosomes in somatic cells is not static but must be maintained by the action of condensin throughout mitosis. In the absence of condensin, chromatin appears to be fully compacted in volume but lacks a normal architecture. Rigidity of mitotic chromosomes is essential for proper kinetochore and microtubule attachments and also for correct shaping and positioning of the mitotic spindle. Wild-type levels of condensin are particularly critical to ensure the completion of sister chromatid separation in anaphase even though the bulk of sister chromatid resolution normally occurs during prophase (Nagasaka et al., 2016).

## MATERIALS AND METHODS

### Cell culture, transfection, siRNA

Chicken DT40 (B lymphoma) cells were cultured in RPMI1640 medium supplemented with 10 % fetal bovine serum and 1% chicken serum at 39°C in 5% CO_2_ in air.

Stable transfection of DT40 cells was performed as described previously (Samejima et al., 2008). We utilized Neon setting 5 (ThermoFisher Scientific) to transiently transfect plasmid DNA (4-10 μg) or siRNA oligonucleotides (2 μl of 100 μM) into 2-4 million cells suspended in 100 μl buffer R from the Neon kit. MAD2 siRNA duplex oligonucleotides were synthesized: (5’-GAAAGCCAUUCAGGAUGAAAUUCGA-3’) and (5’-UCGAAUUUCAUCCUGAAUGGCUUUC-3’) (ThermoFisher Scientific). Control oligonucleotide (AllStars Negative Control siRNA) was purchased from QIAGEN.

### Cell lines

*SMC2 conditional DT40 knockout (SMC2^ON/OFF^) cells:* SMC2^ON/OFF^ cells were previously established (Hudson et al., 2003). In brief, the endogenous *SMC2* gene locus (exon 1-6 including the start codon) was disrupted by Histidinol resistance cassette. These cells depend on the expression of a *GgSMC2* cDNA driven by a tetracycline suppressible promoter (SMC2^ON^). Addition of doxycycline stops the expression of SMC2 protein. SMC2 protein was depleted in the cells treated with doxycycline for > 30 h (SMC2^OFF^).

*SMC2*-*AID*-*GFP DT40 cells:* SMC2-AID-GFP cells were based on the SMC2^OFF^ cells which were constantly cultured in the presence of doxycycline (0.5 μg/ml) to so that non-tagged GgSMC2 protein was not present in the cells. GgSMC2 cDNA fused with minimal AID (mAID) tag (AtIAA17^65-132^) (Natsume et al., 2016) and GFP tag (SMC2-mAID-GFP) at the C-terminus was driven by a 3.8 kb fragment of the GgSMC2 promoter (Hudson et al., 2003). SMC2-mAID-GFP was present as the sole source of SMC2 protein in SMC2-AID-GFP cells (Figure 1A). A CMV promoter drove the expression of plant specific F-box protein OsTIR1 linked to a MmDHFR cDNA by a T2A peptide (synthesized at Thermofisher Scientific and cloned into pCDNA3). 10 μM Methotrexate (MTX) was used to select cells expressing MmDHFR as well as OsTIR1 at a high level. Plasmids coding SMC2-mAID-GFP and OsTIR1 were randomly integrated into the genome.

*CDK1^as^ DT40 cells:* CMV promoter drove the expression of the XICdk1^as^ cDNA (Hochegger et al., 2007) linked to a puromycin resistance gene by a T2A peptide. The GgCdk1 gene was inactivated by transient transfection of plasmids encoding hCas9 cDNA (addgene #41815) and a GgCdk1 guideRNA (based on addgene #41824). The target sequence of the guide RNA was AAAATACGTCTAGAAAGTG.

Where desired, a CENP-H-GFP targeting vector (gift of Yasunari Takami, University of Miyazaki) was used to tag the endogenous CENP-H locus in DT40 cell lines. PACT-RFP construct is a gift of Viji Draviam (University of Cambridge).

### Antibodies and drug treatments

Antibodies used for immunoblotting and indirect immunofluorescence analysis were: rabbit anti-SMC2 (1:500) (Saitoh et al., 1994), guinea pig anti-Ggtopo 2α (1:1000) (Earnshaw et al., 1985), rabbit anti-GgCENP-T (1:1,000) (Hori et al., 2008b), and rabbit anti-MAD2 (1: 100 gift of T. Fukagawa), rabbit-anti GgcyclinB2 (gift of Eric Nigg), mouse anti-α-tubulin B512 or DM1A (1:1,000-4,000; Sigma-Aldrich), mouse anti-Cdk1 monoclonal A17 (1: 200-500: Abcam), and rabbit anti-histone H3 phospho Ser 10 (D2C8) (1:1600, Cell Signaling).

Nocodazole dissolved in DMSO was added to a final concentration of 0.5 μg/ml (Sigma-Aldrich). Doxycycline dissolved in water was added to a final concentration of 0.5 μg/ml (BD) unless stated otherwise. 1NMPP1 dissolved in DMSO was added to a final concentration of 2 μM. Indole-3-acetic acid (auxin) dissolved in ethanol was added to a final concentration of 125 μM unless stated otherwise (Fluka).

### Use of CDK1as system

Typically CDK1^as^ cells were treated with 2 μM 1NMPP1 for 4 h to accumulate cells in G_2_ phase. 1NMPP1 is a bulky ATP analogue that reversibly and specifically inhibits analogue sensitive kinase mutants. 1NMPP1 treatment longer than 4 h results in an increase of multipolar spindles due to centrosome amplification during the G_2_ block. However, CDK1^as^ cells blocked at G_2_ phase for more than 10 h could still enter mitosis synchronously after 1NMPP1 washout without any apparent problems apart from the multipolar spindle. In order to release from the 1NMPP1 block, cells were washed two to three times with fresh media.

### Indirect immunofluorescence of fixed cells

Cells were fixed in pre-warmed 4 % Formaldehyde/PBS for 10 min and permeabilized with 0.15 % triton for 5-10 min. Cells were blocked with 5 % BSA/PBS for 30 min. 1^st^ antibody diluted in the blocking buffer was applied to the cells for 1 h. Cells were washed with PBS 3x for 5 min. 2^nd^ antibody diluted in the blocking buffer (1:500-1:1000, Alexa Fluor 488, 555 or 594, 647 (Molecular Probes, ThermoFisher Scientific) was applied to the cells for 30 min. Cells were washed with PBS for 5 min three times. DNA was stained with Hochest33452 and mounted with Prolong diamond (Molecular Probes, ThermoFisher Scientific).

#### Delta Vision microscopy

3D datasets were acquired using a cooled CCD camera (CoolSNAP HQ; Photometrics) on a wide-field microscope (DeltaVision Spectris; Applied Precision) with a 100× NA 1.4 Plan Apochromat lens. The datasets were deconvolved with softWoRx (Applied Precision), converted to Quick Projections in softWoRx, exported as TIFF files, and imported into Adobe Photoshop for final presentation.

#### 3D structured illumination microscopy (3D-SIM)

Super-resolution images were acquired using structured illumination microscopy. Samples were prepared on high precision cover-glasses (Zeiss, Germany). 3D SIM images were acquired on a N-SIM (Nikon Instruments, UK) using a 100x 1.49NA lens and refractive index matched immersion oil (Nikon Instruments). Samples were imaged using a Nikon Plan Apo TIRF objective (NA 1.49, oil immersion) and an Andor DU-897X-5254 camera using 405, 488 and 561nm laser lines. Step size for Z stacks was set to 0.120 μm as required by manufacturers software. For each focal plane, 15 images (5 phases, 3 angles) were captured with the NIS-Elements software. SIM image processing, reconstruction and analysis were carried out using the N-SIM module of the NIS-Element Advanced Research software. Images were checked for artefacts using the SIMcheck software (http://www.micron.ox.ac.uk/software/SIMCheck.php). Images were reconstructed using NiS Elements software (Nikon Instruments) from a z stack comprising at least 1 μm of optical sections. In all SIM image reconstructions the Wiener and Apodization filter parameters were kept constant. 3D datasets were visualized and analysed using Imaris V8.4 (Bitplane, Oxford Instruments, Oxfordshire, UK).

#### Zeiss Airyscan microscopy

Images of chromosome scaffold proteins on chromosome spreads were obtained using the Airyscan module on a Zeiss LSM 880 confocal, using a x100 alpha Plan-Apochromat objective. Hoechst 33342 was detected using a 405 nm Diode laser and a 420-480 nm bandpass emission filter, Alexa488 was detected using the 488 nm line of an Argon laser and a 495-550 nm bandpass emission filter, the Alexa555 was detected using a HeNe561 laser and a 570 nm long pass emission filter. Alexa647 was detected using a 633 laser and a 605 nm long pass emission filter. Step size for Z stacks was set to 0.145 μm. 3D datasets were visualized and analysed using Imaris V8.4 (Bitplane, Oxford Instruments, Oxfordshire, UK).

### Live cell imaging with Zeiss Airy microscopy

Cells were centrifuged and suspended into Leibovitz’s L-15 Medium (Thermo Fisher Scientific) supplemented with 10% FBS and 1% Chicken serum. DNA was visualised by 1 μM SiR DNA (Spirochromome). GFP was detected using the 488 nm line of an Argon laser and a 495-550 nm bandpass emission filter, RFP was detected using a HeNe561 laser and a 570 nm long pass emission filter. SiR DNA was detected using a 633 laser and a 605 nm long pass emission filter. Step size for Z stacks was set to 0.4 μm. Images were taken every 5 min. 3D datasets were visualized and analysed using Imaris V8.4 (Bitplane, Oxford Instruments, Oxfordshire, UK).

### Intrinsic mitotic chromosome structure (IMCS) assay

Cells were enriched in mitosis either by 12h nocodazole treatment (0.5 μg/ml) (Figure 6A-C) or by 1NMPP1 treatment (Figure 8A, B, D). Auxin was added at the indicated time points (Figure 8A). The cells were plated on polylysine-coated slides for 30 min before the following treatments. The slides were rinsed with PBS, immersed in TEEN buffer for 5 min (1 mM triethanolamine/HCl, pH 8.5, 0.2 mM NaEDTA, and 25 mM NaCl), before immersion into RSB buffer for 15 min (10 mM Tris/HCl, pH 7.4,10 mM NaCl, and 5 mM MgCl_2_). Following two cycles of TEEN buffer/RBS buffer treatments, the cells were fixed with 4% Formaldehyde in RBS buffer and mounted with Vectashield (containing DAPI). The final chromosome morphologies were classified as reformed (green), partially reformed (yellow) or disorganised (magenta).

### Quantitative immunoblotting

Membranes were incubated with the relevant primary antibodies recognizing α-tubulin (as a loading control), SMC2, and MAD2, then subsequently with IRDye-labeled secondary antibodies (LI-COR Biosciences). Fluorescence intensities were determined using a CCD scanner (Odyssey; LI-COR Biosciences) according to the manufacturer’s instructions.

### Flow cytometry analysis

GFP-positive and -negative living cells were analyzed using a FACSCalibur flow cytometer following the manufacturer’s instructions.

DNA contents: Cells were fixed with ice-cold 70% ethanol for overnight. These cells were rinsed with PBS then resuspended with PBS containing 100 μg/ml RNAse A and 5 μg/ml Propidium iodide and were analyzed using a FACSCalibur flow cytometer following the manufacturer’s instructions.

### Measurement of the angle of chromosome mass and centrosomes

SMC2-AID-GFP/CDK1^as^ cells expressing PACT-RFP was treated with 1NMPP1 for 4 h with/without a 3 h auxin treatment. 60 min after 1NMPP1 washout, cells were fixed in pre-warmed 4 % Formaldehyde/PBS for 10 min. DNA was stained with Hochest33452 and mounted with Prolong diamond (Molecular Probes, ThermoFisher Scientific). Pictures of late prometaphase and metaphase cells were taken using Zeiss Airy microscopy as described above. (X, Y, Z) coordinates of the centre of homogenous mass of PACT-RFP signals (X_1_, Y_1_, Z_1_) and (X_3_, Y_3_, Z_3_), and chromosomes (X_2_, Y_2_, Z_2_) were obtained using Imaris V8.4 (Bitplane, Oxford Instruments, Oxfordshire, UK). The angle of chromosome mass and centrosomes was calculated based on the coordinates.

## SBF-SEM

### Preparation of cells

Cells were seeded onto gridded dishes (MatTek) and fixed with 3 % glutaraldehyde and 1% paraformaldehyde in 0.1 M sodium cacodylate buffer for 1 h at RT. Cells were then washed with PBS 3 × 5 min and samples prepared for SBF SEM (West et al., 2010). Extra contrasting steps were introduced, compared to those used for standard TEM to reduce charging and improve the signal to noise ratio. In detail, following fixation, the cells were post fixed and stained with reduced osmium, (2 % osmium tetroxide in dH_2_O + 1.5 % Potassium Ferrocyanide in 0.1M sodium cacodylate buffer) for 1 h at RT. This was followed by 0.1 % tannic acid in ddH_2_O for 20 min at RT. A second osmication step (2 % in ddH_2_O for 40 min at RT), preceded an overnight incubation in aqueous 1 % uranyl acetate at 4°C. The next day cells were stained with Walton’s lead aspartate (0.02 M in lead nitrate + 0.03 M in aspartic acid in ddH_2_O, adjusted to pH 5.5) for 30 min at 60°C. To prevent precipitation artefacts the cells were washed for a minimum of 5 x 3 min with ddH_2_O between each of the staining steps described. Next, samples were dehydrated in a graded ethanol series of 30 %, 50 %, 70 %, 90 % in ddH_2_O for 5 min each, followed by 2 x 5 min 100% ethanol. Samples were then infiltrated with TAAB Hard Premix resin at ratios of 1:1, 2:1 and 3:1 with resin: 100 % ethanol, 30 min per incubation. Finally, samples were incubated in 100 % resin for 2 x 30 min, before embedding the whole dish in 2 mm of 100 % fresh resin. Samples were cured for 48 h at 60°C.

### Preparation of blocks for 3View SBF-SEM

Resin is separated from the gridded dish by trimming away the excess plastic and carefully sliding a razor between the dish and the resin (Booth et al., 2013). Excess resin is removed using a junior hacksaw and scalpel before the block is mounted onto a cryo-pin, cell side up using superglue. Targeted trimming is performed using an ultra-microtome and etched coordinates (Booth et al., 2013).

### SBF-SEM Imaging and Acquisition

Samples were painted with Electrodag silver paint (avoiding the block face) and then coated with 10nm AuPd using a Q150T sputter coater (Quorum Technologies). The sample was inserted into the Gatan 3View sample holder and adjusted so the block face would be central in the microtome and parallel with the knife-edge. Consecutive sectioning and imagining was performed. To obtain a typical resolution of 12 nm in x and y, a frame width of 1024 x 1024 was used. Section thickness was 60 nm over 200-400 sections.

### 3D reconstruction, modeling and segmentation

3View EM stacks were annotated using Amira (FEI). Chromosomes present in every orthoslice were annotated using *masking* and *thresholding* alone (fully automated) or in combination with *magic wand* and *blow* tools (semi automated).

The modelled complement of chromosomes was segmented into discernable isolated objects using *interactive thresholding* and *separate objects* modules. Objects were separated using 3D interpretation and a neighbourhood criteria of 26 connected elements, by at least one corner, edge or face. The marker contrast range (*H*-*extrema*) was set between 5 and 7, depending on the sample. *Label analysis* modules were used to measure the geometry of all isolated structures. Surface renders were generated using unconstrained smoothing at levels 5-7.

### 1NMPP1 Analogue Synthesis

Synthesis of the ATP analog 4-amino-1-tert-butyl-3-(1’-naphthylymethyl)pyrazolo[3,4-d] pyrimidine (1NM-PP1) was carried out in five steps as described by (Hanefeld et al., 1996) with the following modifications. The starting material was 1-naphthaleneacetic acid and the first two steps were carried out under N_2_. In Step 2, triethylamine was used in place of NaH and the product 1-naphthaleneacetyl malonylnitrile was recrystallized from acetonitrile and chloroform. The enol ether product of Step 3 was purified on a silica gel column (elution with 3:1 hexane:ethyl acetate) and stored under N_2_. The final product, a white solid that gave a single spot on TLC, was purified similarly and further recrystallized from ethyl acetate. Its identity was confirmed by ^1^H NMR, HH COSY and high resolution mass spectrometry. ^1^H NMR (270 MHz, CDCl_3_) gave peaks at 1.84 (singlet, 9H), 4.75 (singlet, 2H), 4.85 (broad singlet, 2H), 7.18 (doublet, 1H), 7.38 (triplet, 1H), 7.54 (multiplet, 2H), 7.79-7.92 (multiplet, 2H), 8.22 (doublet, 1H) and 8.24 (singlet, 1H), in close agreement with published data (Bishop et al., 1999). Mass spectrometry gave a mass of 331.181712 for the molecular ion (calculated mass for C_20_H_21_N_5_, 331.179696). The solid compound was stored at 4°C. Aliquots were dissolved in DMSO at 25 mM and stored at ⍰20°C.

## ACKNOWLEDGMENTS

This work was funded by The Wellcome, of which W.C.E. is a Principal Research Fellow (Grant 107022). The Wellcome Centre for Cell Biology is supported by core funding from the Wellcome [203149]. M.T.K. was supported by JSPS KAKENHI Grants (No. 16K15095) and the JST PRESTO program. Sorting of GFP negative cells was performed in the Flow Cytometry facility, Institute of Immunology & Infection Research, Edinburgh with assistance of Dr. Martin Waterfall. We thank Itaru Samejima for comments on the manuscript.

## Author contributions

KS, experimental design, data acquisition, analysis and interpretation, wrote the manuscript; DB electron microscopy, manuscript revision; HO, MP and CW helped to make cell lines. JP and LX synthesized 1NMPP1; MK contribution of new experimental tools and manuscript revision; WCE, experimental design, data analysis and interpretation, co-wrote the manuscript.

## Figure supplement

**Figure 1-figure supplement 1.**
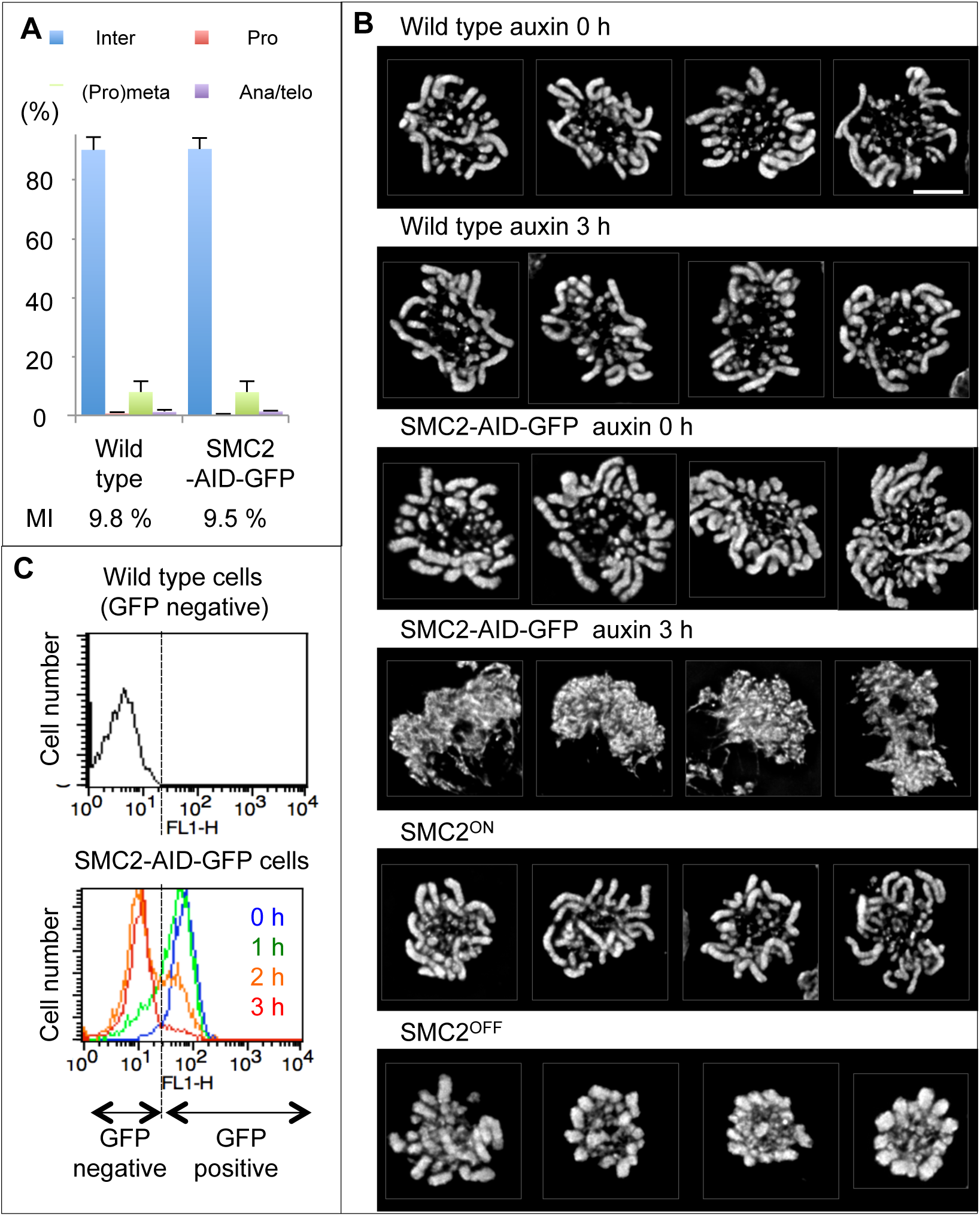
Characterisation of SMC2-AID-GFP cell line. **(A)** Asynchronous wild type/CDK1^as^cells and SMC2-AID-GFP/ CDK1^as^ cells were fixed with formaldehyde and stained with DNA (blue), α-tubulin (green) and H3S10ph (red). 500 cells were counted for each sample. Mean and SD from three independent experiments are shown. **(B)** Cells were treated as Figure 1G. Scale bar 5 μm. **(C)** Flow cytometry analysis of GFP positive cells treated with auxin for 0-3 h.

**Figure 2-flgure supplement 1.**
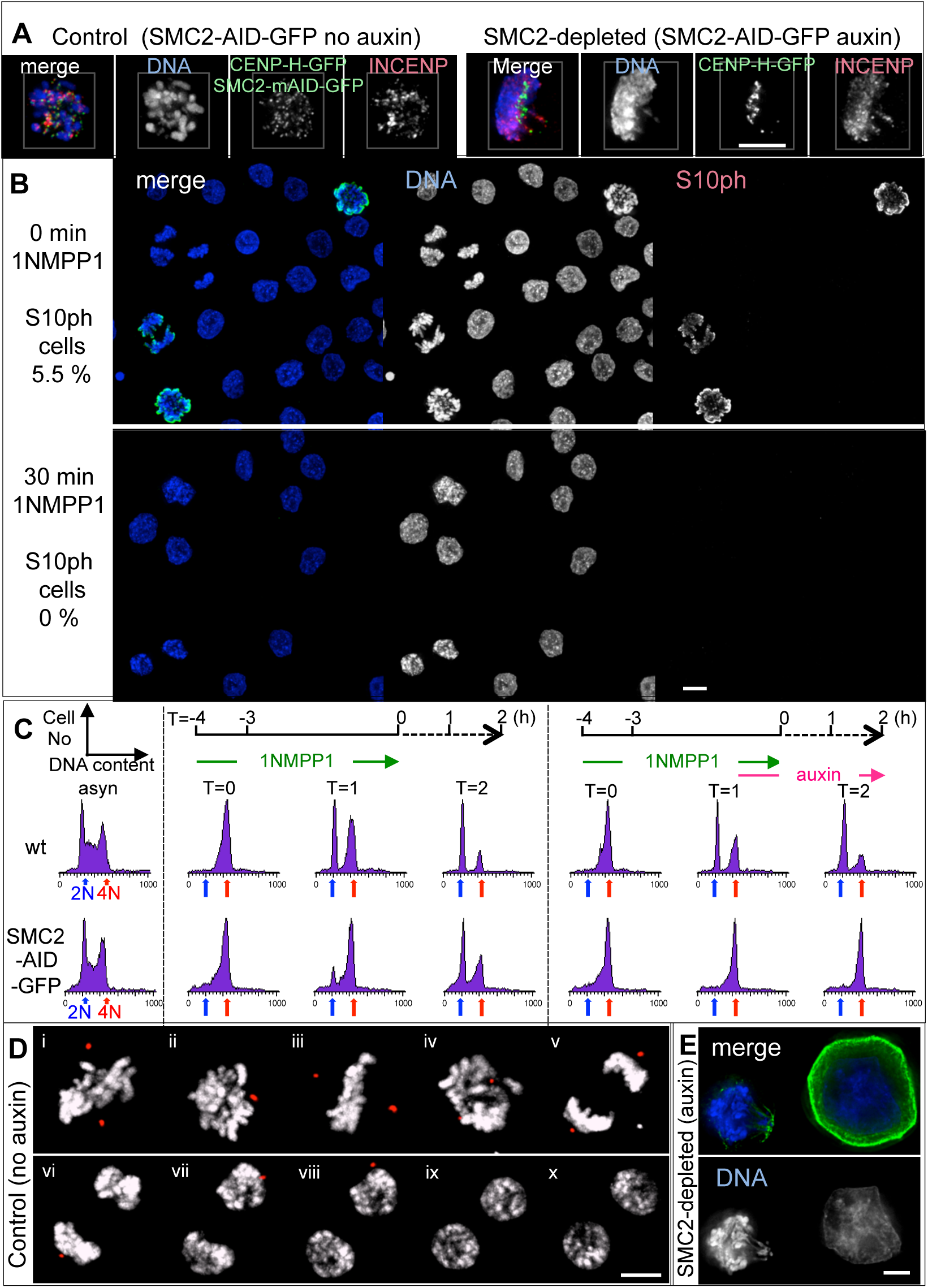
Mitotic cells depleted of SMC2. **(A)** INCENP (red) localises on chromosomes in early mitosis in SMC2-AID-GFP/CDK1^as^/CENP-H-GFP cells both treated or not treated with auxin. Control (no auxin) and SMC2-depleted (auxin 3 h). **(B)** Asynchronous wild type/CDK1^as^ cells were treated with 1NMPP1 for 0 or 30 min and stained for DNA (blue) and H3S10ph (red). **(C)** Analysis of DNA content of wild type/CDK1^as^ cells (wt) and SMC2-AID-GFP/CDK1^as^ cells using flow cytometry. Cells from asynchronously growing culture (i), 4 h block and release from 1NMPP1 (ii), or 4 h block and release from 1NMPP1 with auxin (iii) were analysed as described in Materials and Methods. Positions corresponding cells with 2N DNA (blue arrow) or 4N DNA (red arrow) are shown. **(D)** Stills from live-cell imaging of SMC2-AID-GFP/CDK1^as^ expressing Pericentrin/AKAP450 centrosomal targeting (PACT)-RFP (control). DNA was stained with SiR-DNA. 3D image stacks were collected at 0.4-μm z increments on a Zeiss Airy microscope every 5 mins. SMC2-AID-GFP/CDK1^as^ cells exited mitosis (v-x). Stills of an SMC2-depleted cell (auxin) is shown in Figure 2E. **(E)** SMC2-AID-GFP cells treated with auxin for 2 days. Scale bars 5 μm.

**Figure 3-movie supplement.**

3D reconstruction of control (1) and SMC2-depleted cells (2) shown in Figure 3A taken by 3D-SIM microscopy. DNA (blue), CENP-H-GFP (green), α-tubulin (red).

**Figure 4-figure supplement 1.**
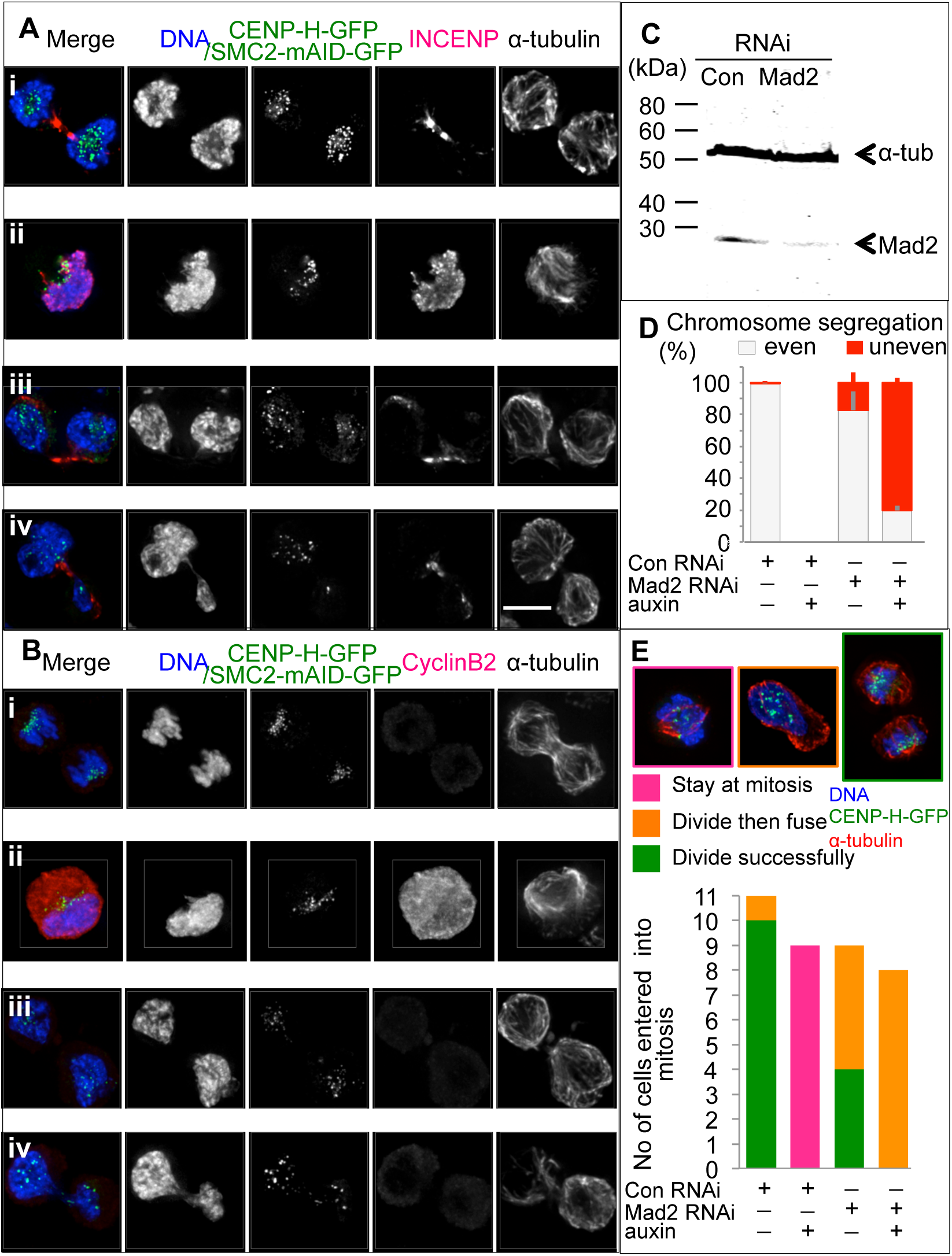
SMC2-depleted cells exit mitosis with highly uneven chromosome segregation after Mad2 RNAi. **(A)** INCENP concentrates on central spindle in cells exiting mitosis. Separate tubulin image is shown. Panels show (i) no auxin, control siRNA; (ii) auxin, control siRNA; (iii) no auxin, Mad2 siRNA; (iv) auxin, Mad2 siRNA. DNA (blue), CENP-H-GFP + SMC2mAID-GFP (green), INCENP (red), α-tubulin (grey). Scale bar 5 μm. **(B)** Cyclin B2 levels fall in cells exiting mitosis. Separate tubulin image is shown. Panels show (i) no auxin, control siRNA; (ii) auxin, control siRNA; (iii) no auxin, Mad2 siRNA; (iv) auxin, Mad2 siRNA. DNA (blue), CENP-H-GFP + SMC2-mAID-GFP (green), Cyclin B2 (red), α-tubulin (grey). Scale bar 5 μm. **(C)** Immunoblot analysis showing MAD2 depletion by specific siRNA. **(D)** Even/uneven chromosome segregation was scored for >50 anaphase or telophase cells per sample. Mean and SD from three independent experiments are shown. **(E)** Quantification of live cell imaging. Cells were transiently transfected with Histone H2B-RFP.

**Figure 6-figure supplement 1.**
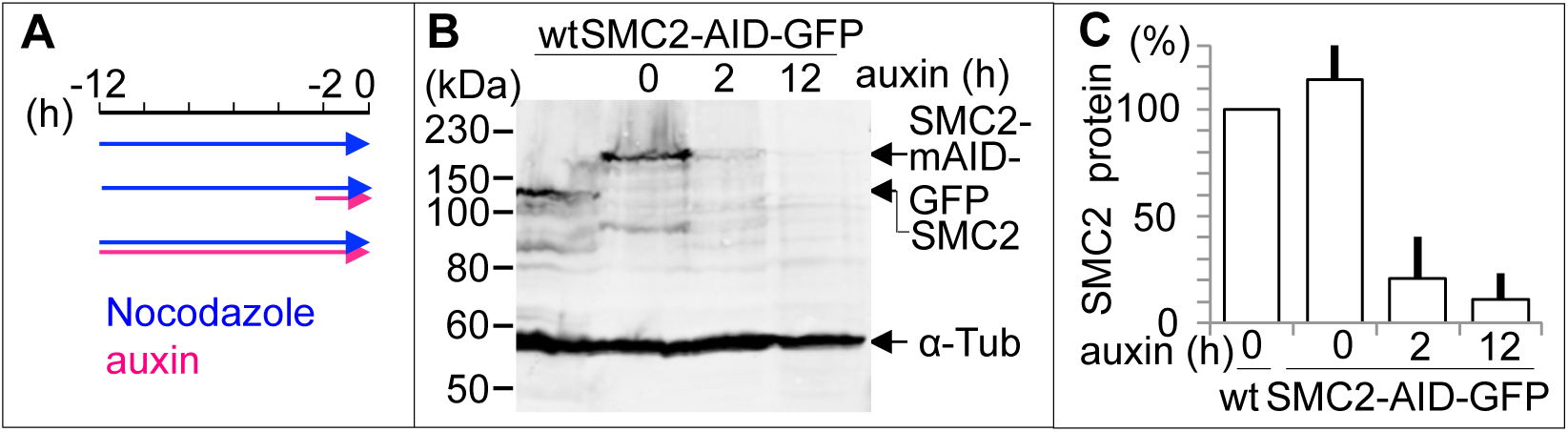
ICMS assay after nocodazole block. **(A)** SMC2-AID-GFP cells were treated with nocodazole for 12 h. Auxin was added at the indicated time points. **(B)** Depletion of SMC2 was confirmed by immunoblot analysis. **(C)** Quantification of SMC2 relative to α-tubulin based on (B). Mean and SD from three independent experiments are shown.

**Figure 6-flgure supplement 2.**
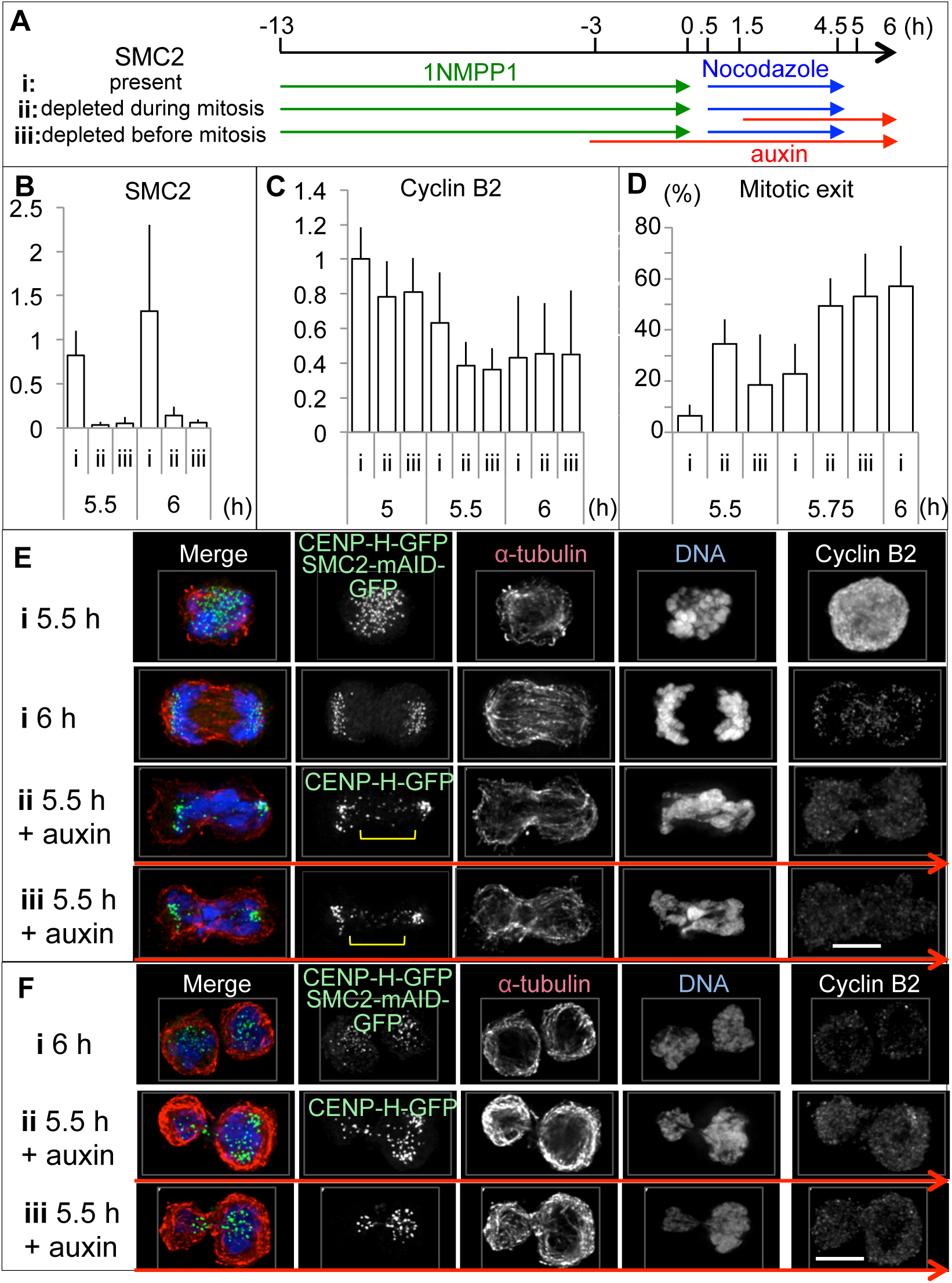
Mitotic exit of cells depleted of SMC2. **(A)** Experimental protocol. **(B)** The relative amount of SMC2 was measured by immunoblot analysis using anti-SMC2 and anti α-tubulin antibodies. **(C)** The relative amount of CyclinB2 was measured by immunoblot analysis using anti-Cyclin B2 and anti α-tubulin antibodies. **(D)** Cells were fixed with 4 % formaldehyde at the time points indicated and stained by DAPI before counting. > 100 cells/sample were counted. For B-D, mean and SD from three independent experiments are shown. **(E)** Cells were fixed with formaldehyde in PBS without hypotonic treatment at the indicated time points and stained with anti-tubulin (red), Cyclin B2 antibodies (grey) and DAPI (blue). CENP-H-GFP + SMC2-mAID-GFP are green. Scale bar 5 μm.

